# Genomic approaches reveal an endemic sub-population of gray wolves in Southern China

**DOI:** 10.1101/512921

**Authors:** Guo-Dong Wang, Ming Zhang, Xuan Wang, Melinda A. Yang, Peng Cao, Feng Liu, Heng Lu, Xiaotian Feng, Pontus Skoglund, Lu Wang, Qiaomei Fu, Ya-Ping Zhang

## Abstract

Despite being one of the most widely distributed terrestrial mammals, the history of gray wolves (*Canis lupus*) in China is not well understood as their habitats have been destroyed with growing economic development. Using six specimens from wolf skins in Chinese Natural History museums, we sequenced their genome using a modified ancient DNA procedure. Using whole genome sequence analysis, we showed that gray wolves from Southern China (SC) derive from a single lineage, distinct from gray wolves from the Tibetan Plateau (*Canis lupus chanco*) and Northern China, suggesting that SC gray wolves may form a distinct sub-population. Of SC gray wolves, one wolf from Zhejiang carries a genetic component from a canid that must have diverged earlier from other wolves than jackals did, perhaps through gene flow from a population related to or further diverged from wolves than the dhole, a species distributed in Southern China and Southeast Asia. This may indicate that interspecific gene flow likely played an important role in shaping the speciation patterns and population structure in the genus *Canis*. Our study is the first to survey museum genomes of gray wolves from Southern China, revealing the presence of an endemic population with ancient interspecific gene flow from a population related to the dhole, and highlighting how sequencing the paleogenome from museum specimens can help us to study extinct species.

## RESULTS AND DISCUSSION

The place of origin for domestic dogs (*Canis lupus familiaris*) remains a controversial question for the scientific community despite many efforts at studying dog domestication [1–7]. Geographic distribution, population structure, and genomic features of wild ancestors are essential factors to determine sources of domestication [8]. Gray wolves (*Canis lupus*) are the closest wild relative of dogs, and they are also one of the most widely distributed terrestrial mammals, originally inhabiting major parts of Eurasia, North America, and North Africa [9–11]. Previous studies suggested gray wolves have a complex history [12, 13], with subpopulation structure related to local niches [14–17] and long-term genetic admixture not only with dogs [5, 18], but also with coyotes [13, 16, 19–21]. In China, gray wolves were distributed across the mainland, including most southern regions [8, 22]. Genomic approaches using gray wolf specimens from Southern China may help to shed new light on the demographic history of gray wolves and domestic dogs.

From two Chinese Natural History museums, we obtained six historical wolf skin samples from gray wolves collected from Mainland China (Figure 1A, S1, and Table 1, detailed description in [8]). Four of them came from Southern China. Two gray wolves were from downstream of Yangtze River of Southern China, collected in Lin’an, Zhejiang province in 1974 and Jiangxi province in May 1974. Another two specimens were collected from Guizhou province near Southeast Asia in 1963. The last two specimens were collected from Northeast China, in Baoqing, Heilongjiang province on Jan 24th, 1957, and Baicheng, Jilin province on Feb 11th, 1956.

**Figure 1.**
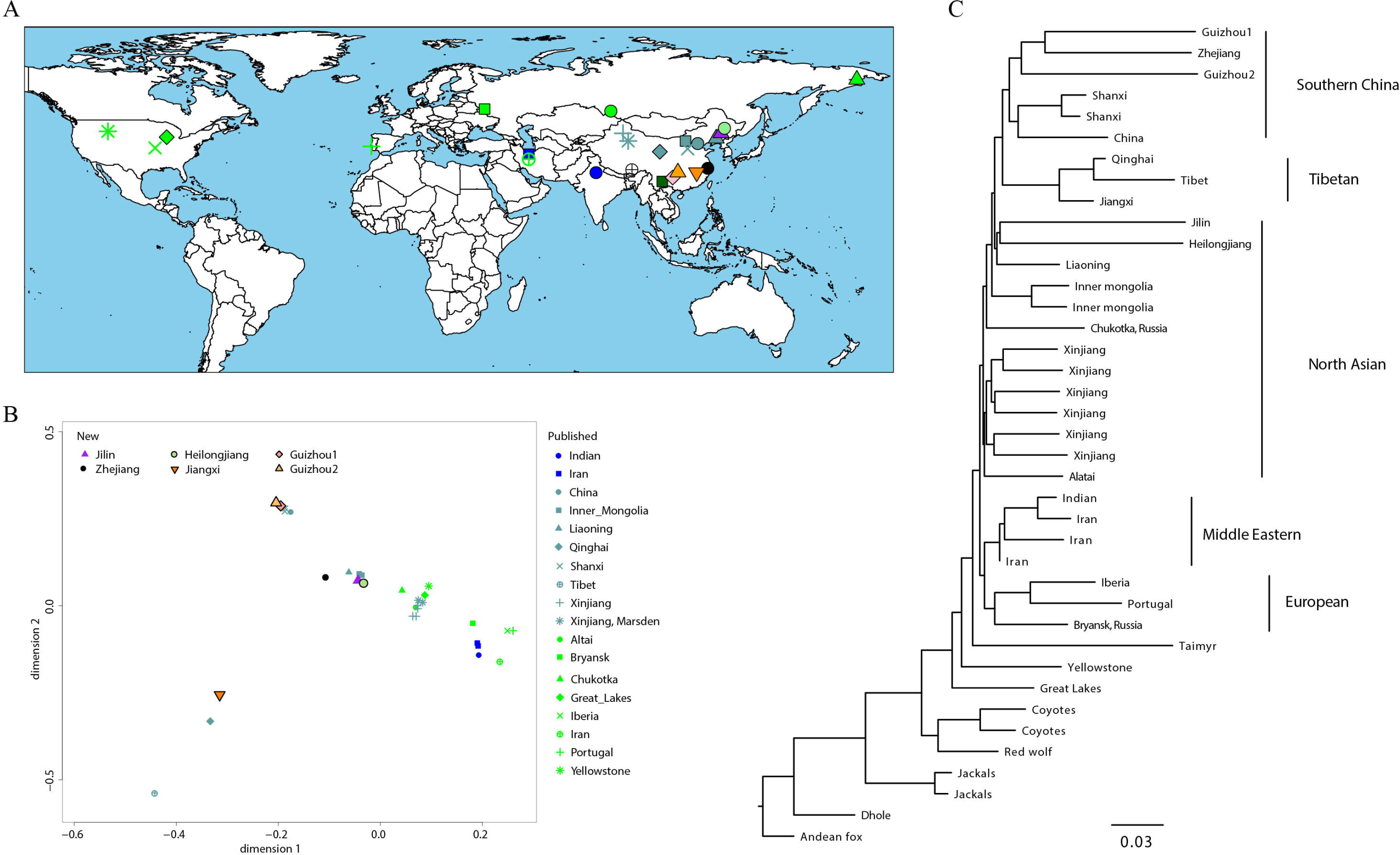
Population structure and phylogeny of 32 gray wolves. (A) Geographic locations, where the key is shared with the principal component analysis in (B). In (B), the label ‘New’ represents the six samples sequenced in the study, and the label ‘Published’ represents 25 samples from previous studies. (C) The maximum likelihood tree of 39 canids, where the Andean fox is used as an outgroup.

**Table 1.** Information on samples in the present study.

| Sample ID | Coverage | Snps | Sources | Location |
| --- | --- | --- | --- | --- |
| Jilin | 1.87 | 6,890,236 | National Zoological<br>Museum of China,<br>Beijing, China | Jilin |
| Zhejiang | 0.15 | 1,866,574 |  | Zhejiang |
| Heilongjiang | 1.61 | 5,438,606 |  | Heilongjiang |
| Guizhou1 | 15.33 | 12,688,043 | Natural History<br>Museum of Zoology,<br>Kunming, China | Guizhou |
| Guizhou2 | 12.39 | 12,968,150 |  | Guizhou |
| Jiangxi | 37.00 | 13,528,667 |  | Jiangxi |

As skin samples were treated with chemicals reagents and underwent special processing for preservation during storage and exhibition in museums, we used a modified ancient DNA (aDNA) protocol [23] to retrieve genetic material from the skin samples. In total, 35 genomic libraries were produced using a double stranded library preparation protocol [24, 25], and each was treated with uracil-DNA-glycosylase (UDG) and endonuclease (Endo VIII) to remove characteristic aDNA deamination [26] (Table S1). We sequenced the libraries using 2×150 bp reads on an Illumina HiSeq X platform.

All but one sample was sequenced to 0.15-15-fold average coverage, too low a coverage to determine heterozygous sites, so we called haploid alleles from a randomly chosen sequence read at each position. We applied a filter where we ignored fragments with length less than 30, ignored the first and last two base pairs of each fragment, and required a base pair quality higher than 20 and mapping quality of at least 30. The Jiangxi wolf was sequenced to 37x (Table 1) – high enough to call heterozygotes. Thus, we applied the software GATK with the Unified Genotyper parameter to determine diploid calls [27]. We also included 79 canids from previous studies [28, 29], including an ancient gray wolf, Taimyr [30], 25 modern gray wolves chosen from regions overlapping the current ranges [3, 7, 18, 19, 31–34], 46 domestic dogs from all over the world [3, 31, 35, 36], two jackals [33], two coyotes [33], one red wolf [19], one dhole [37], and one Andean fox [28] (Figure 1, Table S2).

### Phylogeny and population structure

To investigate the relationship of the newly sampled individuals to wolf and dog populations, we calculated pairwise allele-sharing distances among all pairs of wolf and dog populations. We applied a principal components analysis (PCA) to the resulting pairwise distance matrix using SMARTPCA [38]. The first principal component distinguishes between gray wolf and dog populations, while the second principal component distinguishes between East Asian and European dogs (Figure S2), consistent with previous studies [1, 3, 4]. To obtain increased resolution, we redid the PCA excluding dog populations (Figure 1B). The resulting PCA shows that the two Guizhou wolves cluster with a Chinese wolf (labeled China) from the San Diego Zoo, whose origin is not recorded [18], and a gray wolf from Linfen, Shanxi, China (labeled Shanxi) sampled in 1988, near the border of Southern China [3]. We also find that the Jilin and Heilongjiang wolves cluster with gray wolves from Inner Mongolia and Liaoning. The Jiangxi wolf is closest to the Qinghai wolf, while the Zhejiang wolf is closest to the cluster containing the gray wolves from Inner Mongolia and Liaoning (Figure S1).

Using both a maximum likelihood (ML) tree (Figure 1C) and a neighbor-joining (NJ) tree with only wolves (Figure S3), we further find that the Zhejiang wolf forms a clade with the Guizhou, Shanxi, and China wolves (Figure 1C). We define these gray wolves as the gray wolves from Southern China (SC) and find that they are most closely related to the Qinghai, Tibet, and Jiangxi wolves, where the Tibetan gray wolf (*Canis lupus chanco*) is a gray wolf sub-species that occupies habitats on the Qinghai-Tibet Plateau [39] and adapts to high-altitude environments [32]. Other gray wolves from Northern Asia (NA) form a clade with SC and Tibetan gray wolves relative to Middle Eastern and European gray wolves, which form a distinct clade. The 35,000-year-old wolf from Taimyr Peninsula in northern Siberia joins at the base of the Eurasian wolf phylogeny, and American wolves separate the earliest from other wolves, consistent with [18].

We used TreeMix [40] to investigate the genetic relationship between historical and present-day wolves. TreeMix determines population structure using maximum likelihood trees and allows for both population splits and potential gene flow by using genome-wide allele frequency data and a Gaussian approximation of genetic drift. The maximum likelihood tree (Figure S4) without admixture (m=0) is consistent with previous patterns, where gray wolves from East Asia form three groups: Guizhou and Zhejiang wolves form a clade with Shanxi and China wolves, the Jiangxi wolf forms a clade with Tibet and Qinghai wolves, and Jilin and Heilongjiang wolves, like other North Asian wolves, are outgroup to SC and Tibetan wolves. All three of these groups form a clade relative to non-Asian wolves.

We measured the shared genetic drift between each newly sequenced individual (X) and other dogs and wolves (Y) since their separation from an outgroup, *Dhole*, using *f*_*3*_*(X, Y; Dhole)* [37, 41], and found a similar pattern as above, where the Zhejiang wolf shares the most genetic drift with gray wolves from Guizhou and Jiangxi (Figure S5). We then used *D*-statistics [42] of the form *D(Fox, Test; X, Y)*, where X and Y are previously published wolves, to formally test the relationship these historical wolves have with different wolf populations. For Guizhou and Zhejiang wolves, we find that they share more alleles with the SC gray wolves (Shanxi and China) than with all other wolves, as *D(Fox, Guizhou/Zhejiang; X, SC)*>0 (10.8<Z<33.4, Table S3). The Jiangxi wolf shares more allele with the Tibet and Qinghai gray wolves than other wolves, as *D(Fox, Jiangxi; X, Tibet/Qinghai)*>0 (11.7<Z<24.9, Table S3), while Jilin and Heilongjiang wolves share the most alleles with NA gray wolves. These results support our above analyses, again grouping the Guizhou and Zhejiang wolves with the SC wolves, the Jiangxi wolf with the Tibetan wolves, and the Jilin and Heilongjiang wolves with the NA wolves.

In summary, our results revealed that the lowland Chinese wolves [20] consisted of two major populations: SC and NA wolves. Gray wolves from Zhejiang and Guizhou group most closely with and share the most genetic drift with SC wolves, which includes present-day populations in Shanxi and China. The Jilin and Heilongjiang gray wolves share the most genetic similarity to the NA gray wolves, which is the other clade in Northern China and Eastern Russia.

### Testing for admixture in gray wolves

Using *D(Fox, X; Test, Y)*, we find that all gray wolves (X) share more alleles with the Guizhou, Jilin, and Heilongjiang wolves (Test) than with the gray wolves from Tibet and Qinghai, i.e. *D(Fox, X; Test, Qinghai/Tibet)<0* (Table S5). The Jiangxi wolf shares more alleles with the Tibet and Qinghai wolves than other wolves (Table S3), and here, we find that the Tibet and Qinghai wolves share more alleles with the Jiangxi wolf, i.e. *D(Fox, Qinghai/Tibet; Jiangxi, X)<0* (−28.7<Z<−14.7, Table S5), emphasizing that the Jiangxi, Tibet, and Qinghai wolves form a clade. However, while *D(Fox, X; Jiangxi, Qinghai)~0* (−2.3<Z<1.8, Table S5), indicating a symmetric relationship as expected for the Jiangxi and Qinghai wolves forming a clade, we observe *D(Fox, X; Jiangxi, Tibet)<0* (Table S5), suggesting that the Jiangxi wolf has a connection to other wolves relative to the Tibetan wolf. Thus, we observe that the Tibet and Qinghai gray wolves act as an outgroup to most SC and NA gray wolf populations, and while the Jiangxi wolf forms a clade with the Tibet and Qinghai gray wolves, the Jiangxi wolf shows connections to non-Tibetan wolf populations. To test for admixture between the SC, NA, and Tibetan wolves, we used *f3(Test; X, Y)*, where a significantly negative value (|Z|>3) suggests that the Test population is a mixture of ancestry related to X and Y, two other wolf populations. Testing all combinations as both a source population and the admixed population, we found that *f3(Jiangxi/Qinghai; Tibetan, SC/NA)<0* (−19.101<Z<−9.141, Table S4), suggesting that both the Jiangxi and Qinghai wolves show evidence of ancestry from populations related to both Tibetan and SC gray wolves, explaining why the Jiangxi gray wolf shares a connection to non-Tibetan gray wolf populations.

In contrast to all other ancient gray wolves from China, the Zhejiang wolf shows a markedly different pattern, where all other wolf populations, including the Tibetan and Qinghai gray wolves and more distantly related wolves from further west, form a clade with each other relative to the Zhejiang wolf. That is, we observe that *D(Fox, X; Zhejiang, Y)>0* (9.1<Z<71.5, Table S5), where X and Y are all other gray wolves, including the Taimyr. Earlier, we found that *D(Fox, Zhejiang; X, Shanxi/China)>0* (Table S3), indicating that the Zhejiang wolf shows connections to the wolves from Shanxi and China. These results suggest that the Zhejiang wolf shows a close relationship to gray wolves from Shanxi and China, but that this wolf also possesses an ancestral component that is older than the common ancestral population of the Taimyr and all other gray wolves. The error rate for the Zhejiang wolf (0.4%) is higher than that estimated for other wolves sequenced in this study (0.1%-0.2%), likely because of its low coverage (Table 1). After simulating an error rate similar to that observed for the Zhejiang wolf in these other wolves, we find that our results remain consistent with our previous results. That is, the Zhejiang wolf shows a distinct pattern from that observed in other wolves for both lower and higher error rates.

We use the genomic data from canids typically outgroup to all wolves and dogs – the Dhole, Jackal, Coyote, and Red wolf – to understand how deeply the old component found in the Zhejiang wolf separated from other canid populations. Other canids separated from wolf populations very early, with the Dhole diverging earliest, followed by the Jackal and most recently the Coyote [36, 43, 44]. The Red wolf is genetically very similar to the Coyote and shows substantial gene flow from gray wolves [13, 19]. First, comparing the Jackal to wolves (X) and the Coyote, we find that for all wolves but the Zhejiang wolf, *D(Fox, Jackal; X, Coyote)* ranges from −0.04 to −0.02 (−20.2<Z<−12.6, Table S6), indicating a connection between the Jackal and gray wolves. We find the reverse for the Zhejiang wolf, however, where the Jackal shares more alleles with the Coyote than with the Zhejiang wolf, i.e. *D(Fox, Jackal; Zhejiang, Coyote)=0.13* (Z=39.3, Table S6). We find similar results replacing the Coyote with the Red wolf. The large contrast between the results for the Zhejiang wolf compared to other gray wolves suggests that the old component came from a population that diverged deeply in the past, who separated prior to the common ancestor of jackals and coyotes.

We also observe that for all gray wolves, we find *D(Fox, Dhole; X, Jackal)*>0 (Table S6), suggesting that gray wolves share a deep lineage older than the separation of the Jackal and Dhole or that there is a direct genetic connection between the Jackal and Dhole. However, while D ranges from 0.1 to 0.2 (4.5<Z<7.9, Table S6) for most gray wolves, using the Zhejiang wolf greatly increases the D value to 0.11 (Z=22.4, Table S6). We find that *D(Fox, Dhole; Zhejiang, Jackal)* remains significantly positive (Z=12.2) using only transversions, suggesting that the result for the Zhejiang wolf is not related to ancient DNA damage and likely reflects an unusual admixture history. If the Zhejiang wolf was no different from other gray wolves, especially the two Guizhou individuals to which they share the closest relationship (Table S5), we would expect to find that *D(Fox, Dhole; X, Zhejiang)~0*, which would indicate that the Zhejiang wolf and other gray wolves are similarly related to the Dhole. However, we observe that *D(Fox, Dhole; X, Zhejiang*)<<0 (−23.4<Z<−12.9, Table S6), indicating that the Dhole shares more alleles with other gray wolves than with the Zhejiang gray wolf. These patterns suggest that the ancestral component found in the Zhejiang wolf came from a population that diverged earlier than the common ancestor of the Jackal and Dhole, which is older than the separation of the Jackal and Dhole from wolves, suggesting that the Zhejiang wolf possesses very deep ancestry not found in other gray wolves.

Using the tree model with no admixture from Treemix (Figure S6), we visualized the matrix of residuals (Figure S7) to determine how the estimated genetic relationship between each pair of canids fit the model. A high residual indicates that the pair does not fit the tree model and may be candidates for an admixture event. We find two candidate admixture events, the first between the Andean fox and the Zhejiang wolf (Figure 2); and the second between gray wolves from Portugal and Iberia. In a maximum-likelihood tree allowing two admixture events, admixture from the lineage leading to the Andean fox to the lineage leading to the Zhejiang wolf is included, while gray wolves from Portugal and Iberia are instead grouped into the same cluster (Figure S8). Admixture between the outgroup Andean fox and the Zhejiang wolf supports our conclusions from the D-statistic analysis (Table S5 and Table S6), in which the Zhejiang wolf possesses an ancestral component that came from a population that diverged earlier than the Jackal or Dhole did from wolves. The estimated value of the migration event in the Zhejiang individual is 12.3% ± 0.4% (P<2.2×10^−308^). In the Treemix analysis, we used the Andean fox as an outgroup, whose distance from the included canids would result in weak phylogenetic constraints. In addition, we also used the F4-ratio test to estimate the proportion of this deep ancestry, and since it is older than the separation of the Jackal and Dhole, we used an unrooted phylogeny where the Fox is used as a proxy as the source of the deep ancestry. Thus, we estimate the proportion of ancestry related to the Fox, which is given by:

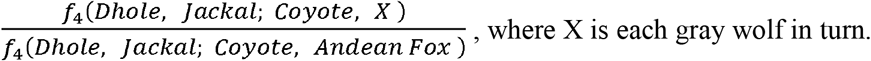

**Figure 2.**
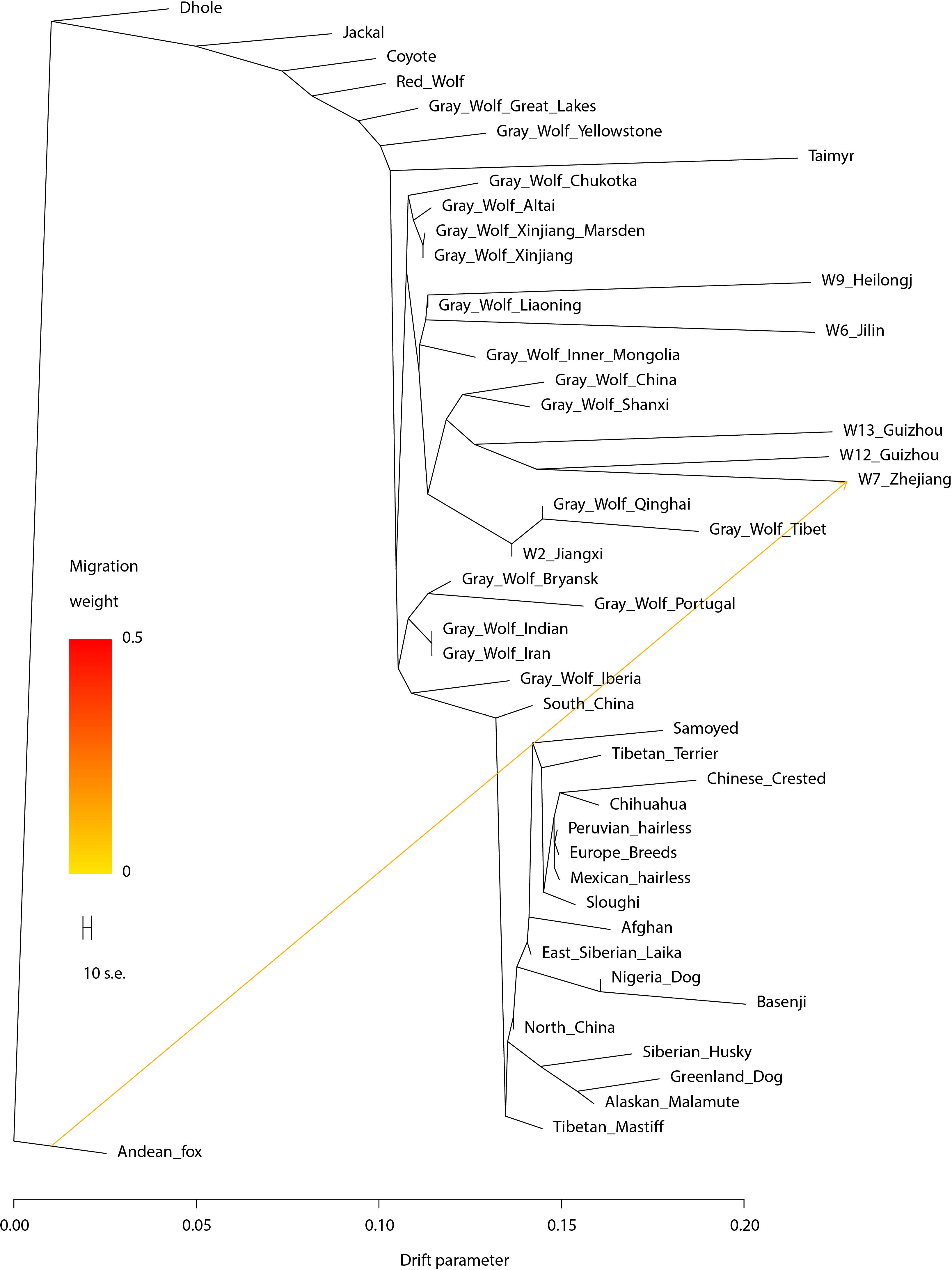
The maximum-likelihood tree based on TreeMix with m=1. The scale bar shows ten times the average standard error of the entries in the sample covariance matrix. We have used the prefix “Gray_Wolf_” for highlighting specimens of gray wolves.

Using this method, we found that the estimated admixture proportion of the deep ancestry for the Zhejiang wolf is 11.7% ± 0.5%, whereas all other grey wolves have an estimated admixture proportion close to zero. Thus, both the Treemix analysis and the F4-ratio test support the presence of gene flow from an ancient canid population into the ancestors of the Zhejiang wolf.

### Conclusion

The distribution of gray wolves in East Asia is controversial since some studies have claimed that gray wolves never existed [45, 46] or are now extinct from Southern China [47], while others sources, especially those based on Chinese literature, stated that they are present across all of mainland China [8]. In this study, we provide the first comparative genomic analysis of gray wolves from East Asia focusing particularly on wolves from Southern China, where some believed no gray wolves were distributed [46, 48]. Previously, Asian wolves could be divided into two populations: Tibetan gray wolves (*Canis lupus chanco*) and Chinese lowland wolves [20, 32]. Here, using ancient genome-wide data, we reconstruct the phylogeny and evaluate the population structure and shared genetic drift between East Asian gray wolves to show that they form three major groups, which we call Southern China, North Asia, and Tibetan, based on their geographic distribution. Interestingly, specimens from SC gray wolves were all collected from 1956 - 1988. Our results highlight that the population in Southern China is endemic, and with the fast growing economic development of China, it is paramount to protect and restore their ecological habitat. Through our study, we also emphasize the value of using paleogenomic approaches to study the numerous museum specimens available [49, 50], and we address the importance of using population genomics to determine current or future conservation efforts.

Finally, our analyses show that admixture played a large role between the different Asian wolves, and we highlight two instances here. First, we find that a wolf as far southeast as Jiangxi province shows evidence of being a mixture of Tibetan wolves and other wolves in China. Second, we traced an unusual admixture event in the Zhejiang wolf. In many analyses, this wolf behaved similarly to other gray wolves in China, particularly those from Southern China. However, tests of admixture suggest that the Zhejiang wolf has gene flow from a canid that diverged earlier than the separation of wolves and jackals. D-statistic analysis suggests this may be from the Dhole, a species distributed in Southern China and Southeast Asia [51], or another canid that separated earlier than the divergence between wolves and the Dhole. Whatever the source of this ancestry, estimates of the admixture proportion from this deeply diverging population are estimated to be ~12%. Our results, taken together with previous research [52], reveal that canids are an ideal system in which to study how gene flow can shape speciation in a genus, and highlight the need for greater study of ancient gray wolf populations.

## ACCESSION NUMBERS

Sequence data for six gray wolf genomes has been submitted to the Genome Sequence Archive (http://gsa.big.ac.cn/) under accession number PRJCA001135.

## SUPPLEMENTAL INFORMATION

Supplemental Information includes Supplemental Experimental Procedures, nine figures, and six tables, which can be found with this article online at XXX.

## AUTHOR CONTRIBUTIONS

Y.-P.Z. and Q.F. designed research. G.-D.W. and Q.F. managed the project. L.W. collected samples. P.C., F.L., M.Z., and X.F. performed DNA experiments. Q.F., X.F., M.Y., and P.S., did the data processing. Q.F., G.-D.W., M.Z., X.W., M.Y., and H.L. analyzed the data. G.-D.W., M.Y., Q.F., and Y.-P.Z. wrote the manuscript.

## ACKNOWLEDGMENTS

We thank Jun Chen and Gexia Qiao from the National Zoological Museum of China, Beijing, China, and Weiwei Li and Song Li from the Natural History Museum of Zoology, Kunming, China for helping to collect samples. We thank the BIG Data Center at the Beijing Institute of Genomics, Chinese Academy of Sciences (CAS) for use of their high-performance computing platform. This work was supported by the Strategic Priority Research Program (B) (XDB13000000) of the CAS, National Natural Science Foundation of China (91731304, 91731303, 41672021, and 41630102). P.S. was supported by the Francis Crick Institute which receives its core funding from Cancer Research UK (FC001595), the UK Medical Research Council (FC001595), and the Wellcome Trust (FC001595). Q.F. is supported by CAS (XDB26000000, XDA19050102, QYZDB-SSW-DQC003), and the Howard Hughes Medical Institute (grant number 55008731). G.D.W. is supported by the Youth Innovation Promotion Association, CAS, and the CAS international collaborating grant proposal (152453KYSB20150002). This work was supported by the Animal Branch of the Germplasm Bank of Wild Species, CAS (the Large Research Infrastructure Funding).

