## Supplementary Figure and Tables for "Genomic approaches reveal an endemic sub-population of gray wolves in Southern China"

Figure S1-9.

Table S1-6.

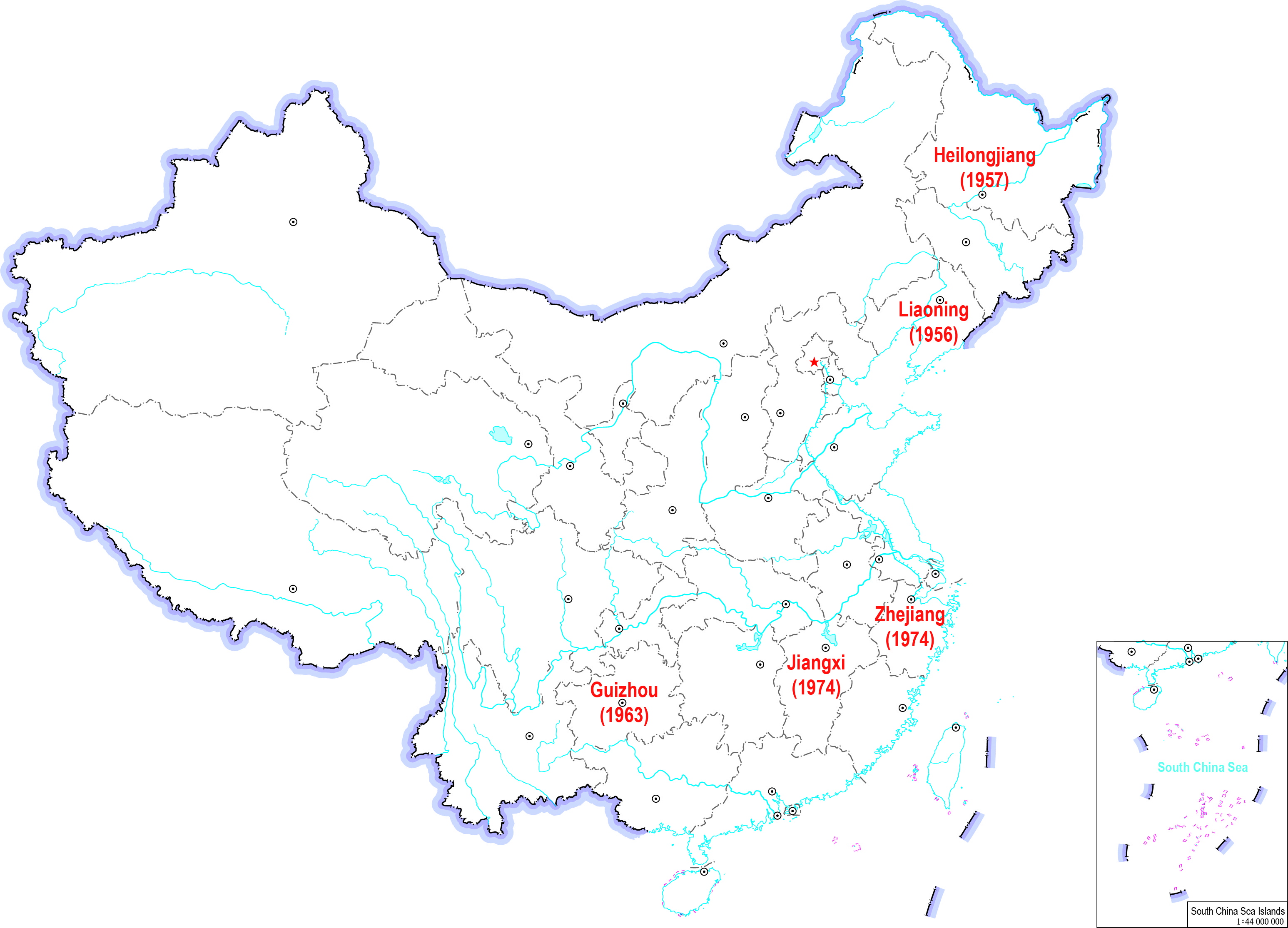

Figure S1. Geographical origin of six museum wolf skin specimens in China. The collected year is indicated within brackets. Guizhou has 2 individuals.

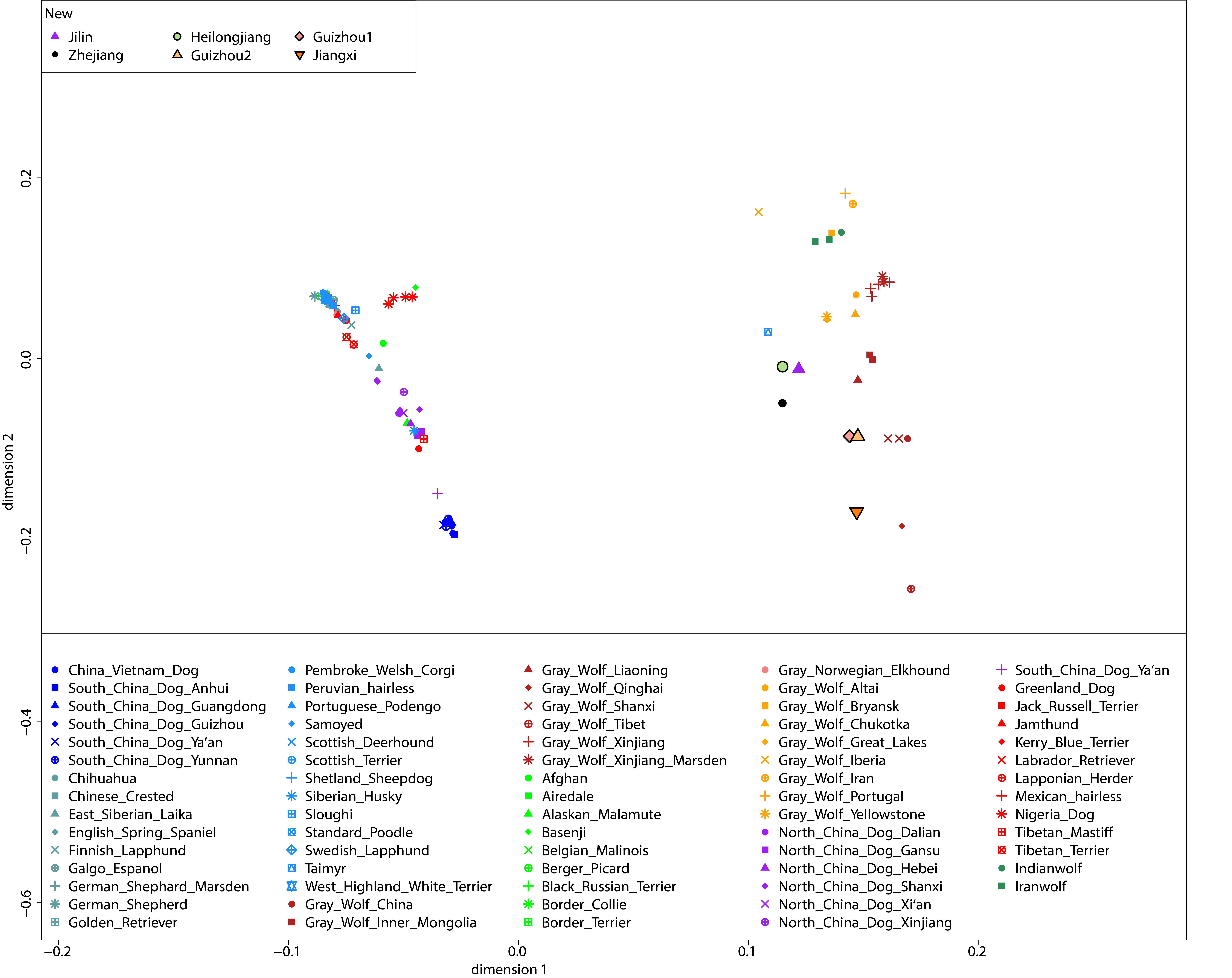

Figure S2. PCA analysis with wolf and dog populations. We have used the prefix "Gray_Wolf_" for highlighting specimens of gray wolves.

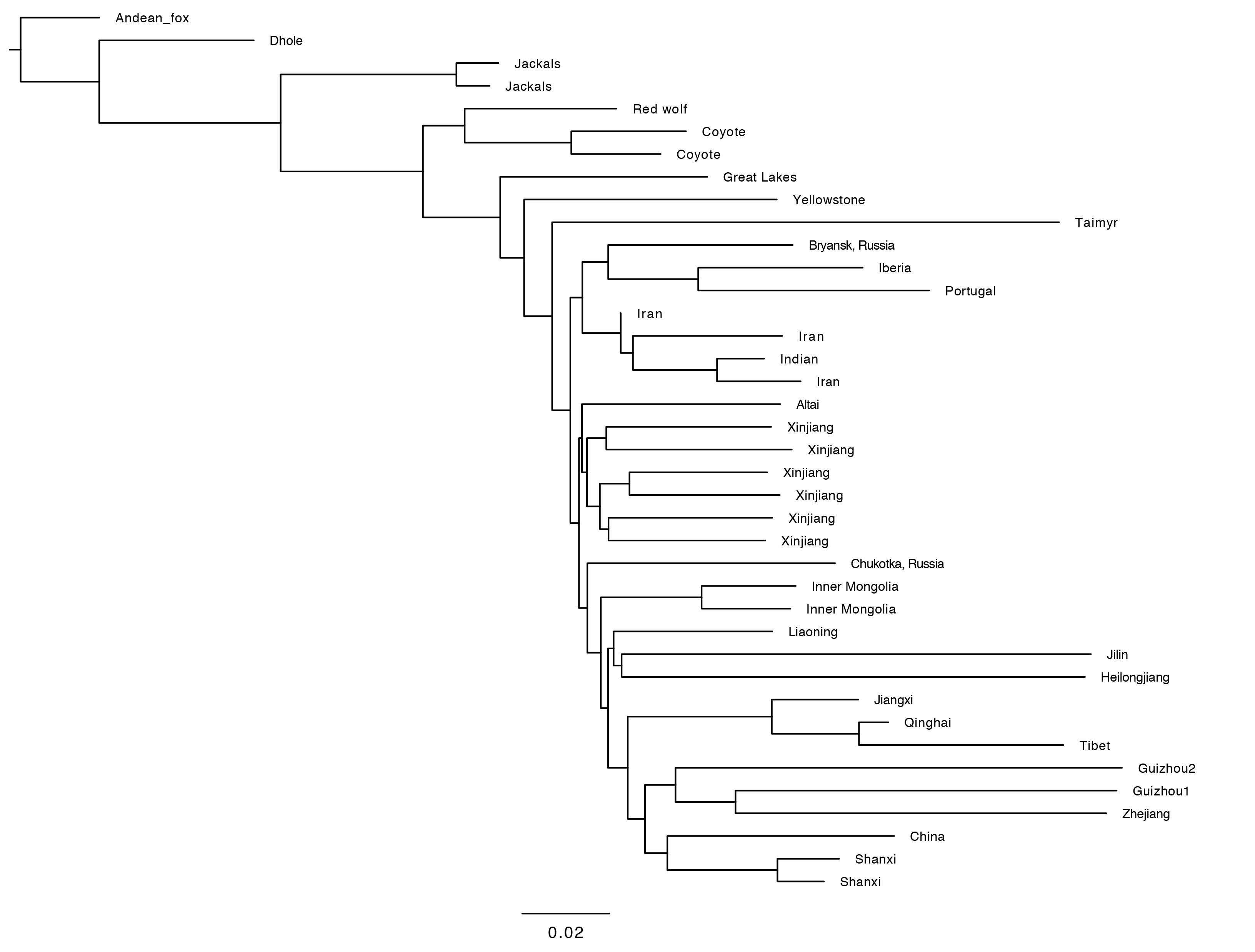

Figure S3. Neighbor-Joining tree including 39 canids without dogs. The Andean fox is the outgroup.

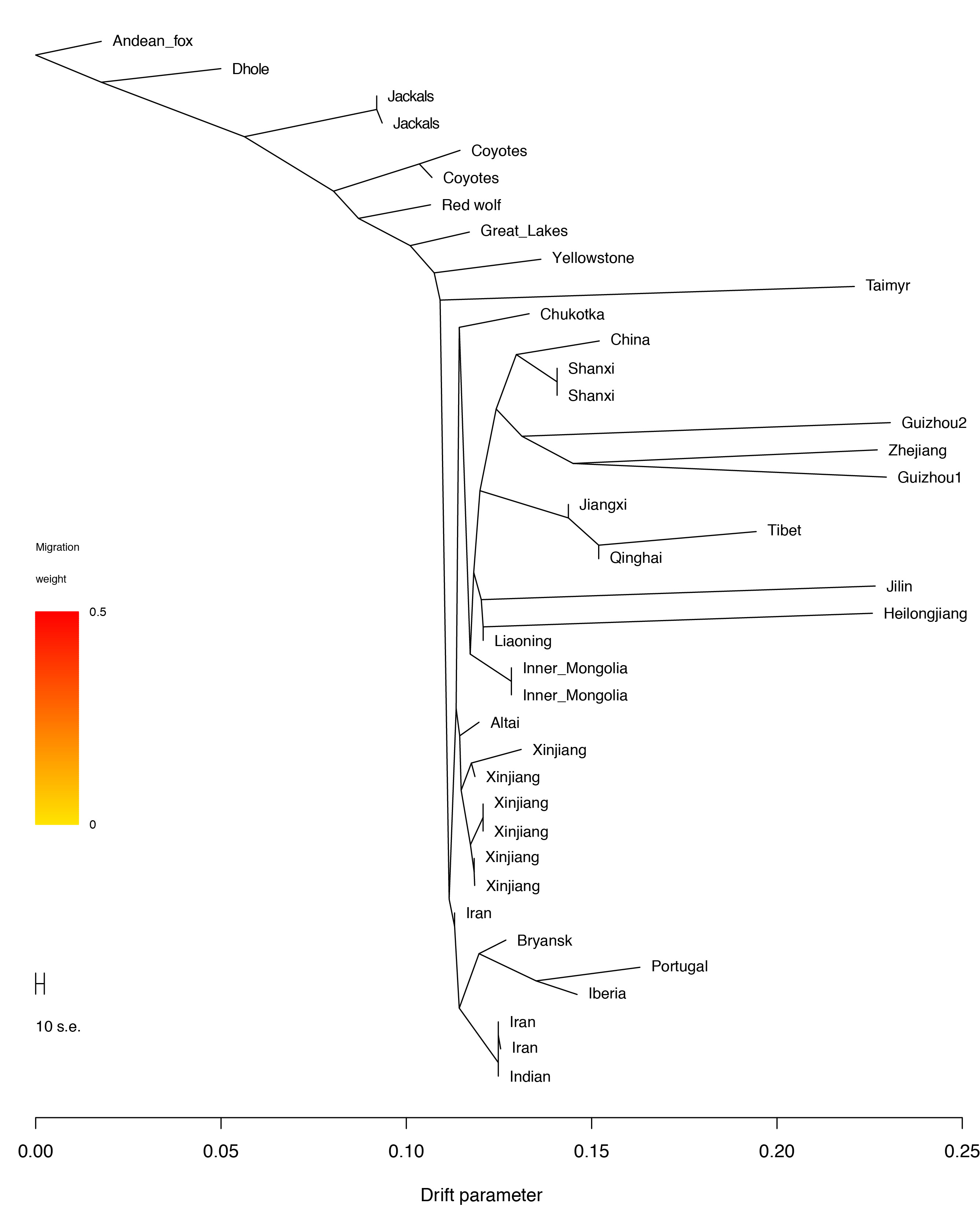

Figure S4. The maximum-likelihood tree based on TreeMix using 39 canids without dogs and m=0. The scale bar shows ten times the average standard error of the entries in the sample covariance matrix.

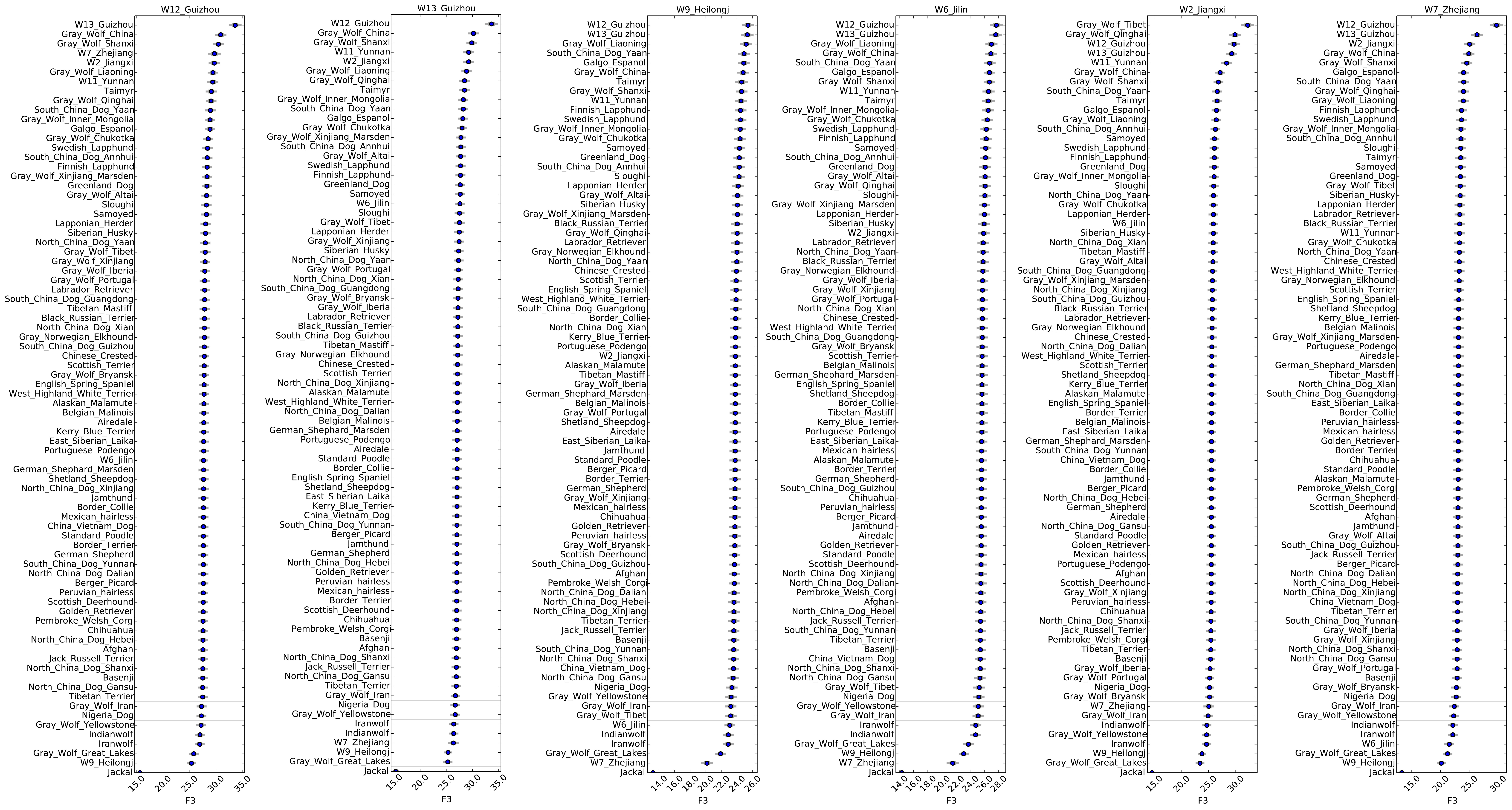

Figure S5. f3(Dhole; X, Y), where X is one of the six newly sequenced wolves and Y are other canids. We have used the prefix "Gray_Wolf_" for highlighting specimens of gray wolves.

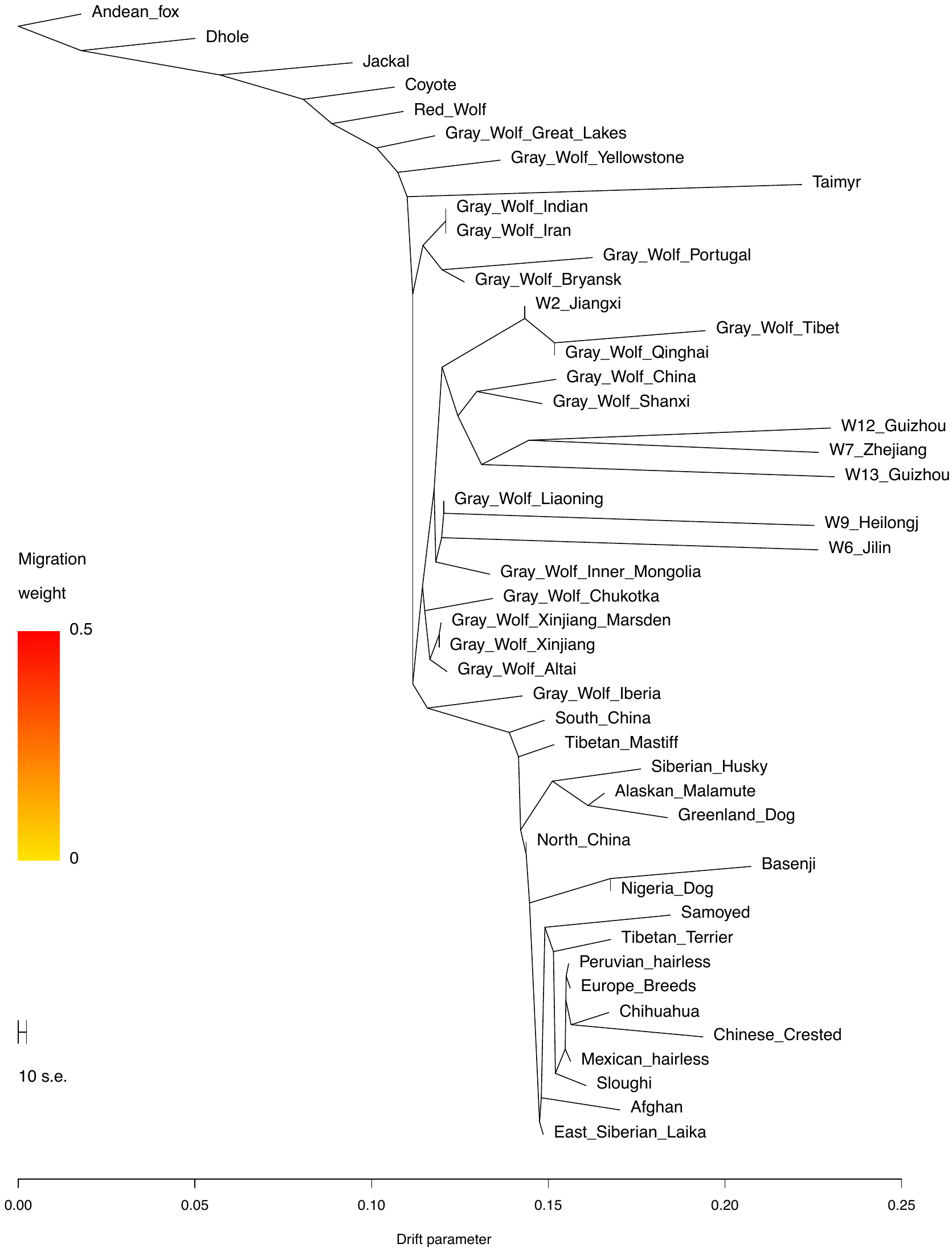

Figure S6. The maximum-likelihood tree based on TreeMix with m=0. The scale bar shows ten times the average standard error of the entries in the sample covariance matrix. We have used the prefix "Gray_Wolf_" for highlighting specimens of gray wolves.

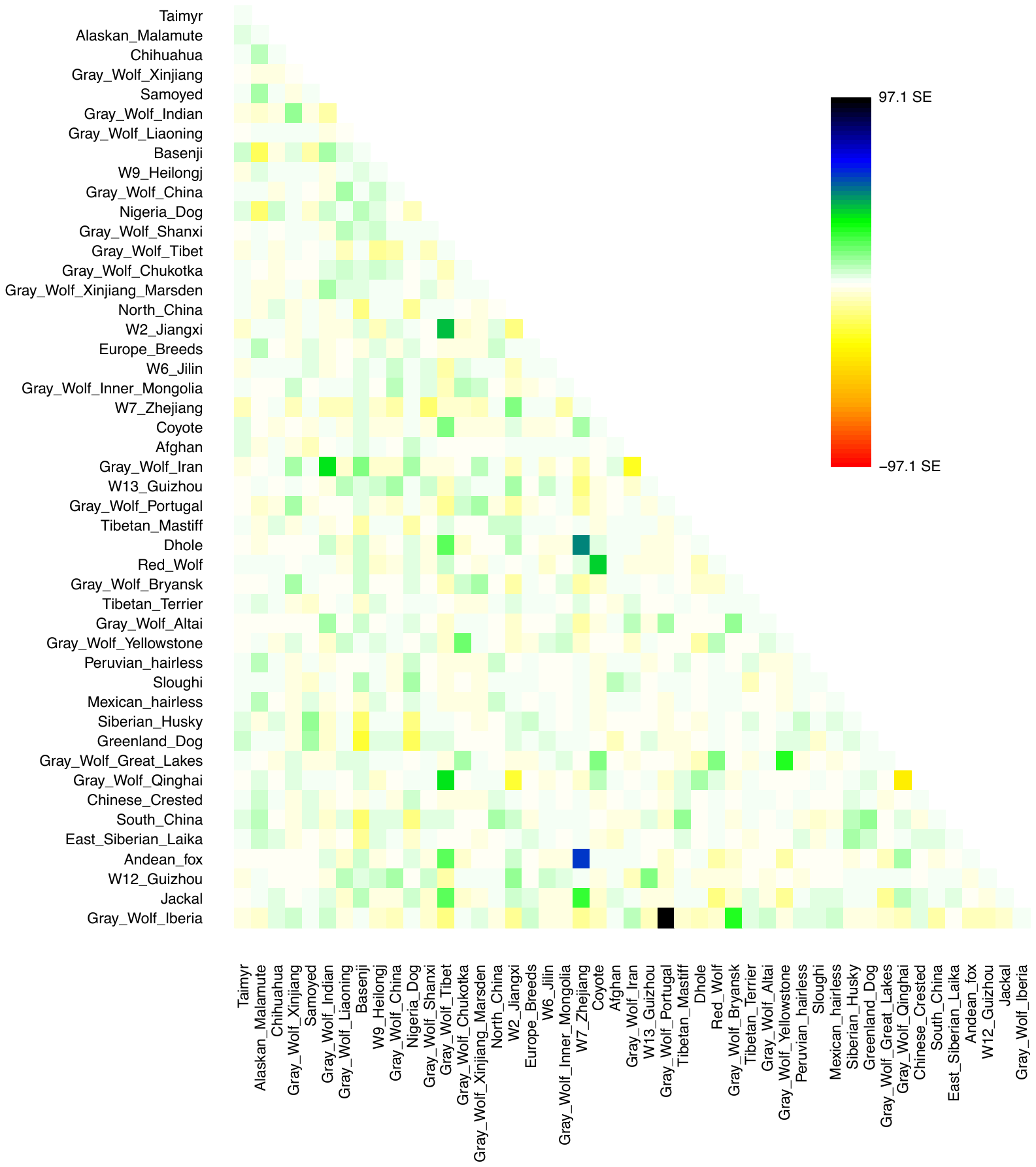

Figure S7. The residual fit from the maximum likelihood tree in Figure S6. This is determined by dividing the residual covariance between each pair of populations by the average standard error across all pairs. Colors are described in the palette on the right. Residuals above zero represent populations that are more closely related to each other in the data than in the best-fit tree, and thus are candidates for admixture events. We have used the prefix "Gray_Wolf_" for highlighting specimens of gray wolves.

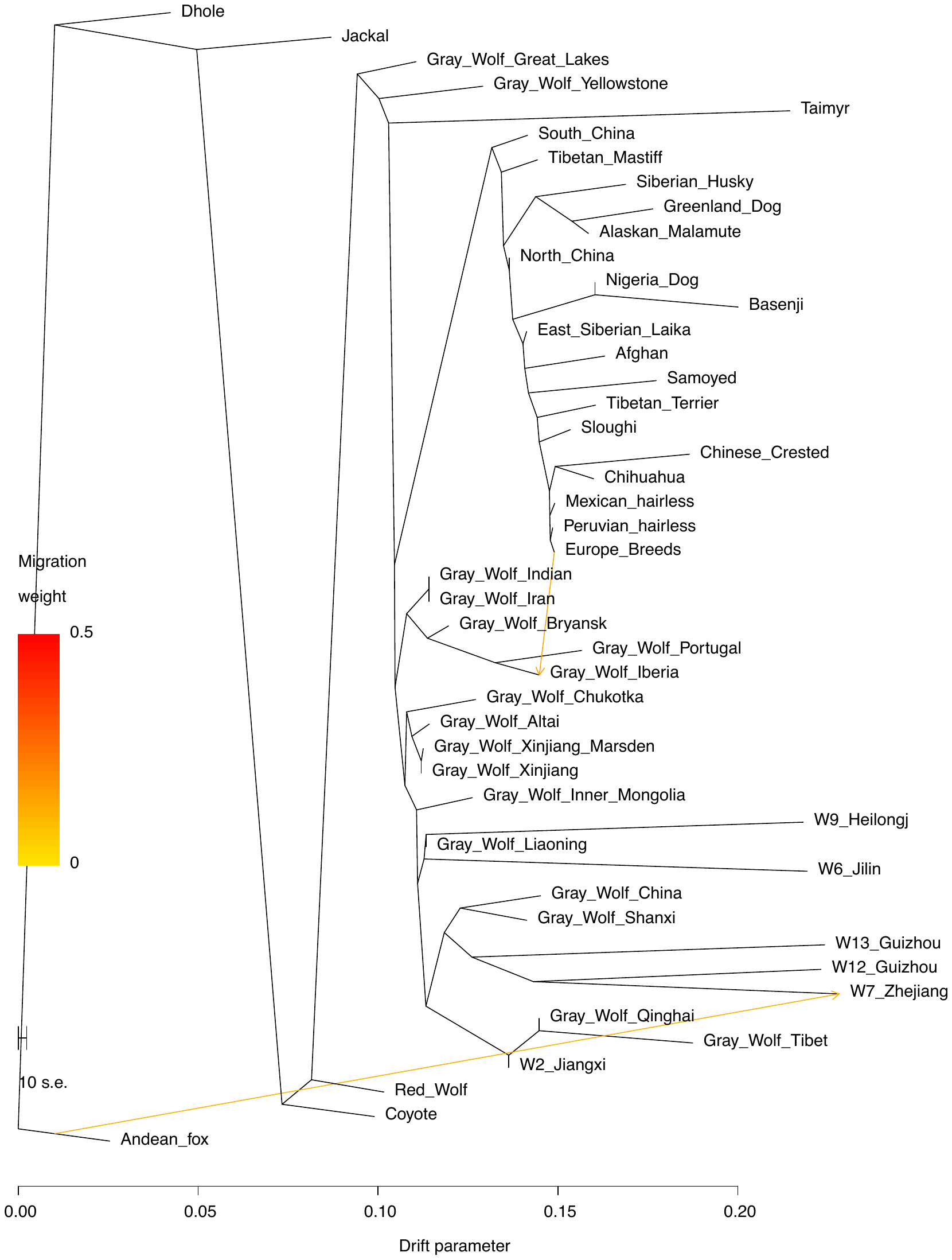

Figure S8. The maximum-likelihood tree based on TreeMix with m=2. The scale bar shows ten times the average standard error of the entries in the sample covariance matrix. We have used the prefix "Gray_Wolf_" for highlighting specimens of gray wolves.

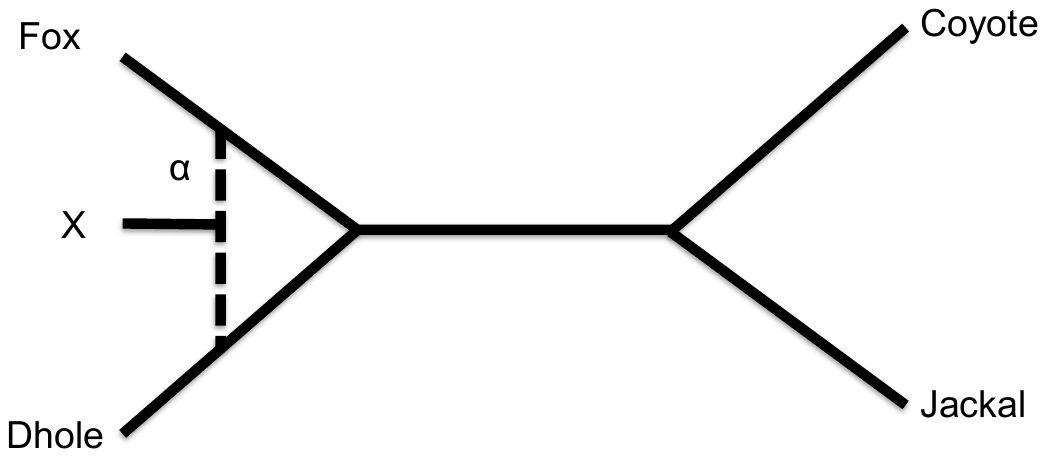

Figure S9. Unrooted tree used to estimate the archaic admixture proportion in the Zhejiang wolf.

Table S1. Sequencing metrics on the libraries of the samples.

| **Sample_ID** | **Lib_ID** | **Raw** | **Merged** | **&L30** | **Mapped** | **%Mapped** | **Unique** | **Average Length** | **Coverage** |
| --- | --- | --- | --- | --- | --- | --- | --- | --- | --- |
| W12 | L4705 | 285343617 | 282720996 | 279349969 | 193639299 | 69 | 87020138 | 66 | 2.280 |
| W12 | L4706 | 122888988 | 121920909 | 120203042 | 81036195 | 67 | 37300930 | 66 | 0.980 |
| W12 | L4723 | 99422832 | 98301650 | 95950384 | 66689079 | 70 | 27296673 | 60 | 0.656 |
| W12 | L4724 | 129941007 | 128741838 | 125878134 | 86419489 | 69 | 35693699 | 60 | 0.859 |
| W12 | L4721 | 458823905 | 457178101 | 448236865 | 354599384 | 79 | 172415148 | 75 | 5.168 |
| W12 | L1236 | 220782482 | 214658250 | 212167619 | 172702204 | 81 | 83601122 | 82 | 2.755 |
| W13 | L4715 | 174435249 | 170656674 | 168612632 | 143289506 | 85 | 67178876 | 71 | 1.912 |
| W13 | L4733 | 100006948 | 97480812 | 96184443 | 79766884 | 83 | 37206113 | 74 | 1.101 |
| W13 | L4716 | 139660264 | 136843871 | 135476042 | 94368328 | 70 | 41458135 | 75 | 1.243 |
| W13 | L4734 | 72759420 | 71413179 | 70557705 | 48841322 | 69 | 20183735 | 72 | 0.585 |
| W13 | L1237 | 376669471 | 358779675 | 356919462 | 259792144 | 73 | 138338276 | 94 | 5.179 |
| W2 | L4717 | 57032184 | 56165346 | 55929745 | 49470031 | 88 | 25470196 | 81 | 0.820 |
| W2 | L4718 | 108657013 | 107230625 | 106675141 | 95573663 | 90 | 49006998 | 78 | 1.537 |
| W2 | L4735 | 84038690 | 82911815 | 82368601 | 73430883 | 89 | 34211885 | 74 | 1.008 |
| W2 | L4736 | 904760479 | 897952829 | 896171857 | 792391018 | 88 | 457142173 | 92 | 16.831 |
| W2 | L1228 | 614619758 | 568519001 | 567900196 | 519132756 | 91 | 335201526 | 126 | 16.915 |
| W6 | L4707 | 127681136 | 126097468 | 118289082 | 90384661 | 76 | 26902055 | 48 | 0.520 |
| W6 | L4708 | 125430513 | 123589141 | 116027856 | 87251813 | 75 | 25879613 | 49 | 0.505 |
| W6 | L4725 | 58051880 | 57192340 | 49981995 | 32368357 | 65 | 10635232 | 43 | 0.183 |
| W6 | L4726 | 77378791 | 76219741 | 65107619 | 47902350 | 74 | 15267977 | 42 | 0.256 |
| W6 | L1232 | 389136153 | 325919319 | 315703233 | 246562564 | 78 | 11324756 | 58 | 0.263 |
| W7 | L4709 | 39112069 | 38593429 | 35402488 | 1843676 | 5 | 732974 | 57 | 0.017 |
| W7 | L4710 | 35581720 | 35083335 | 32713276 | 1193424 | 4 | 395721 | 56 | 0.009 |
| W7 | L4711 | 39015421 | 38438945 | 35639593 | 1833027 | 5 | 705431 | 64 | 0.018 |
| W7 | L4712 | 39853260 | 39242507 | 36562343 | 2739467 | 7 | 1008169 | 63 | 0.026 |
| W7 | L4727 | 39610530 | 39030578 | 32984973 | 2129260 | 6 | 796738 | 47 | 0.015 |
| W7 | L4728 | 35207785 | 34640731 | 29296827 | 1317589 | 4 | 404382 | 46 | 0.007 |
| W7 | L4729 | 44741369 | 44125280 | 37579861 | 2065001 | 5 | 683061 | 55 | 0.015 |
| W7 | L4730 | 35964348 | 35467842 | 30198475 | 2576745 | 9 | 810242 | 54 | 0.017 |
| W7 | L1233 | 216672870 | 196311964 | 188328108 | 8194354 | 4 | 324626 | 89 | 0.012 |
| W9 | L4713 | 60306027 | 59823823 | 56491103 | 48351903 | 86 | 17888738 | 51 | 0.364 |
| W9 | L4714 | 48275546 | 47733287 | 45095180 | 35539238 | 79 | 13910738 | 51 | 0.285 |
| W9 | L4731 | 78659917 | 78002396 | 69178887 | 58004190 | 84 | 20595472 | 44 | 0.359 |
| W9 | L4732 | 67266981 | 66511206 | 60235429 | 47951316 | 80 | 18366379 | 45 | 0.329 |
| W9 | L1234 | 146878870 | 129755983 | 125676741 | 53058660 | 42 | 1949475 | 71 | 0.055 |

Table S2. Genome information from public databases.

| **Species** | **Location** | **ID** | **Data sources** |
| --- | --- | --- | --- |
| **fox** | **Andean** | **Andean_fox** | **Adam Auton et al., 2013, Plos Genetics[1]** |
| **Dhole** | **Beijing Zoo** | **RUFZCHN00001** | **Guo-Dong Wang et al., 2018, National Science Review[2]** |
| **Jackals** | **Krasnodar, Russia** | **AUR008537** | **Xuan Wang et al., 2018, BioRxiv[3]** |
|  | **Krasnodar,Russia** | **AUR008538** | **Xuan Wang et al., 2018, BioRxiv[3]** |
| **Coyotes** | **Monterey area, US** | **LAT007000** | **Xuan Wang et al., 2018, BioRxiv[3]** |
|  | **California, US** | **SAMN02921301** | **Bridgett M. vonHoldt et al., 2016, Science Advances[4]** |
| **Red wolf** | **US** | **SAMN02921317** | **Bridgett M. vonHoldt et al., 2016, Science Advances[4]** |
| **Gray wolves** | **Great Lakes** | **gwglw_RKW2455** | **Bridgett M. vonHoldt et al., 2016, Science Advances[4]** |
|  | **Yellowstone** | **gwynp_RWK1547** | **Bridgett M. vonHoldt et al., 2016, Science Advances[4]** |
|  | **Taimyr** | **Taimyr** | **Pontus Skoglund et al., 2015, Current Biology[5]** |
|  | **Iran** | **LUP004103** | **Xuan Wang et al., 2018, BioRxiv[3]** |
|  | **Iran** | **LUP004107** | **Xuan Wang et al., 2018, BioRxiv[3]** |
|  | **Iran** | **gwirw_RKW3073** | **Clare D. Marsden et al., 2016, PNAS[6]** |
|  | **India** | **SAMN02921311** | **Bridgett M. vonHoldt et al., 2016, Science Advances[4]** |
|  | **Bryansk,Russia** | **LUPWRUS00003** | **Guo-Dong Wang et al., 2013, Nature Communications[7]** |
|  | **Iberia** | **gwibe_XXWIB98** | **Laura R. Botigue et al., 2017, Nature Communications[8]** |
|  | **Portugal** | **gwprt_LOBO423** | **Clare D. Marsden et al., 2016, PNAS[6]** |
|  | **Shanxi** | **LUPZCHN00005** | **Guo-Dong Wang et al., 2015, Cell Research[9]** |
|  | **Shanxi** | **LUPZCHN00006** | **Guo-Dong Wang et al., 2015, Cell Research[9]** |
|  | **China** | **gwcwz_RKW3916** | **Adam H. Freedman et al., 2014, Plos Genetics[10]** |
|  | **Qinghai** | **gwcwq_XinQH11** | **Wenping Zhang, et al., 2014, Plos Genetics[11]** |
|  | **Tibet** | **gwcwt_XinTI09** | **Wenping Zhang, et al., 2014, Plos Genetics[11]** |
|  | **Liaoning** | **LUPWCHN00003** | **Guo-Dong Wang et al., 2015, Cell Research[9]** |
|  | **Inner Mongolia** | **LUPWCHN00001** | **Guo-Dong Wang et al., 2013, Nature Communications[7]** |
|  | **Inner Mongolia** | **LUPZCHN00002** | **Guo-Dong Wang et al., 2015, Cell Research[9]** |
|  | **Chukotka,Russia** | **LUPWRUS00002** | **Guo-Dong Wang et al., 2013, Nature Communications[7]** |
|  | **Xinjiang** | **LUPWCHN00008** | **Guo-Dong Wang et al., 2015, Cell Research[9]** |
|  | **Xinjiang** | **LUPWCHN00009** | **Guo-Dong Wang et al., 2015, Cell Research[9]** |
|  | **Xinjiang** | **LUPWCHN00010** | **Guo-Dong Wang et al., 2015, Cell Research[9]** |
|  | **Xinjiang** | **LUPWCHN00013** | **Guo-Dong Wang et al., 2015, Cell Research[9]** |
|  | **Xinjiang** | **gwcwx_XinXJ24** | **Wenping Zhang, et al., 2014, Plos Genetics[11]** |
|  | **Xinjiang** | **gwcwx_XinXJ30** | **Wenping Zhang, et al., 2014, Plos Genetics[11]** |
|  | **Altai,Russia** | **LUPWRUS00001** | **Guo-Dong Wang et al., 2013, Nature Communications[7]** |
| **Breed dogs** | **Afghan** | **FAMBAFG00001** | **Guo-Dong Wang et al., 2015, Cell Research[9]** |
|  | **Alaskan Malamute** | **FAMBALM00001** | **Guo-Dong Wang et al., 2015, Cell Research[9]** |
|  | **Belgian Malinois** | **FAMBBEM00001** | **Guo-Dong Wang et al., 2015, Cell Research[9]** |
|  | **Chihuahua** | **FAMBCHI00001** | **Guo-Dong Wang et al., 2015, Cell Research[9]** |
|  | **East Siberian Laika** | **FAMBESL00001** | **Guo-Dong Wang et al., 2015, Cell Research[9]** |
|  | **Finnish Lapphund** | **FAMBFIL00001** | **Guo-Dong Wang et al., 2015, Cell Research[9]** |
|  | **Galgo Español** | **FAMBGAL00001** | **Guo-Dong Wang et al., 2015, Cell Research[9]** |
|  | **Gray Norwegian Elkhound** | **FAMBGNE00001** | **Guo-Dong Wang et al., 2015, Cell Research[9]** |
|  | **Greenland dog** | **FAMBGRD00001** | **Guo-Dong Wang et al., 2015, Cell Research[9]** |
|  | **German Shepherd Dog** | **FAMBGSD00001** | **Guo-Dong Wang et al., 2015, Cell Research[9]** |
|  | **Jämthund** | **FAMBJAM00001** | **Guo-Dong Wang et al., 2015, Cell Research[9]** |
|  | **Lapponian Herder** | **FAMBLAH00001** | **Guo-Dong Wang et al., 2015, Cell Research[9]** |
|  | **Mexican naked** | **FAMBMEN00001** | **Guo-Dong Wang et al., 2015, Cell Research[9]** |
|  | **Peruvian naked** | **FAMBPEN00001** | **Guo-Dong Wang et al., 2015, Cell Research[9]** |
|  | **Samoyed** | **FAMBSAM00001** | **Guo-Dong Wang et al., 2015, Cell Research[9]** |
|  | **Siberian Husky** | **FAMBSIH00001** | **Guo-Dong Wang et al., 2015, Cell Research[9]** |
|  | **Sloughi** | **FAMBSLO00001** | **Guo-Dong Wang et al., 2015, Cell Research[9]** |
|  | **Swedish Lapphund** | **FAMBSWL00001** | **Guo-Dong Wang et al., 2015, Cell Research[9]** |
|  | **Tibetan Mastiff** | **FAMBTIM00001** | **Guo-Dong Wang et al., 2015, Cell Research[9]** |
| **Village dogs** | **Xi’an** | **FAMICHN00001** | **Guo-Dong Wang et al., 2013, Nature Communications[7]** |
|  | **Ya’an** | **FAMICHN00003** | **Guo-Dong Wang et al., 2013, Nature Communications[7]** |
|  | **Dalian** | **FAMICHN00004** | **Guo-Dong Wang et al., 2015, Cell Research[9]** |
|  | **Gansu** | **FAMICHN00005** | **Guo-Dong Wang et al., 2015, Cell Research[9]** |
|  | **Gansu** | **FAMICHN00006** | **Guo-Dong Wang et al., 2015, Cell Research[9]** |
|  | **Gansu** | **FAMICHN00007** | **Guo-Dong Wang et al., 2015, Cell Research[9]** |
|  | **Hebei** | **FAMICHN00012** | **Guo-Dong Wang et al., 2015, Cell Research[9]** |
|  | **Shanxi** | **FAMICHN00014** | **Guo-Dong Wang et al., 2015, Cell Research[9]** |
|  | **Shanxi** | **FAMICHN00015** | **Guo-Dong Wang et al., 2015, Cell Research[9]** |
|  | **Shaanxi** | **FAMICHN00016** | **Guo-Dong Wang et al., 2015, Cell Research[9]** |
|  | **Shaanxi** | **FAMICHN00017** | **Guo-Dong Wang et al., 2015, Cell Research[9]** |
|  | **Xinjiang** | **FAMICHN00019** | **Guo-Dong Wang et al., 2015, Cell Research[9]** |
|  | **Ya’an** | **FAMICHN00002** | **Guo-Dong Wang et al., 2013, Nature Communications[7]** |
|  | **Guangdong** | **FAMICHN00010** | **Guo-Dong Wang et al., 2015, Cell Research[9]** |
|  | **Guizhou** | **FAMICHN00011** | **Guo-Dong Wang et al., 2015, Cell Research[9]** |
|  | **Yunnan** | **FAMICHN00021** | **Guo-Dong Wang et al., 2015, Cell Research[9]** |
|  | **Yunnan** | **FAMICHN00023** | **Guo-Dong Wang et al., 2015, Cell Research[9]** |
|  | **Anhui** | **FAMICHN00025** | **Guo-Dong Wang et al., 2015, Cell Research[9]** |
|  | **Ibadan, Nigeria** | **FAMINGR00001** | **Guo-Dong Wang et al., 2015, Cell Research[9]** |
|  | **Ondo, Nigeria** | **FAMINGR00002** | **Guo-Dong Wang et al., 2015, Cell Research[9]** |
|  | **Uyo, Nigeria** | **FAMINGR00003** | **Guo-Dong Wang et al., 2015, Cell Research[9]** |
|  | **Taraba State, Nigeria** | **FAMINGR00004** | **Guo-Dong Wang et al., 2015, Cell Research[9]** |
|  | **China/Vietnam border** | **FAMIVNM00001** | **Guo-Dong Wang et al., 2015, Cell Research[9]** |
|  | **China/Vietnam border** | **FAMIVNM00002** | **Guo-Dong Wang et al., 2015, Cell Research[9]** |
|  | **China/Vietnam border** | **FAMIVNM00003** | **Guo-Dong Wang et al., 2015, Cell Research[9]** |
|  | **China/Vietnam border** | **FAMIVNM00004** | **Guo-Dong Wang et al., 2015, Cell Research[9]** |
|  | **China/Vietnam border** | **FAMIVNM00005** | **Guo-Dong Wang et al., 2015, Cell Research[9]** |

Table S3 Z-score of *D(Fox, test; X, Y).*

| **D(O Guizhou1;P3 P4)** | | | |  |  |  |  |  |  |  |  |  |  |  |  |
| --- | --- | --- | --- | --- | --- | --- | --- | --- | --- | --- | --- | --- | --- | --- | --- |
| **P3/P4** | **Taimyr** | **Iranwolf** | **Indianwolf** | **Portugal** | **Inner_Mongolia** | **Liaoning** | **Xinjiang** | **Altai** | **Chukotka** | **Bryansk** | **Shanxi** | **China** | **Xinjiang_Marsden** | **Qinghai** | **Tibet** |
| **Taimyr** |  | -0.3 | 0.0 | **5.8** | **25.1** | **23.5** | **17.2** | **15.9** | **18.4** | **7.5** | **30.1** | **26.2** | **18.2** | **14.2** | 2.4 |
| **Iranwolf** | 0.3 |  | 0.5 | **7.8** | **28.9** | **25.2** | **22.1** | **17.5** | **21.6** | **9.4** | **33.4** | **26.7** | **21.8** | **15.1** | **2.7** |
| **Indianwolf** | 0.0 | -0.5 |  | **6.3** | **25.0** | **21.9** | **15.2** | **14.4** | **17.0** | **7.1** | **28.5** | **25.2** | **16.4** | **14.8** | 2.5 |
| **Portugal** | **-5.8** | **-7.8** | **-6.3** |  | **19.1** | **18.5** | **9.9** | **9.2** | **12.0** | 1.1 | **27.3** | **23.0** | **11.8** | **10.1** | -0.5 |
| **Inner_Mongolia** | **-25.1** | **-28.9** | **-25.0** | **-19.1** |  | **3.8** | **-15.4** | **-12.7** | **-8.4** | **-20.3** | **14.9** | **13.5** | **-12.0** | **-3.2** | **-11.6** |
| **Liaoning** | **-23.5** | **-25.2** | **-21.9** | **-18.5** | **-3.8** |  | **-15.1** | **-13.1** | **-10.3** | **-17.7** | **10.8** | **11.2** | **-13.2** | **-5.4** | **-13.1** |
| **Xinjiang** | **-17.2** | **-22.1** | **-15.2** | **-9.9** | **15.4** | **15.1** |  | 1.1 | **5.0** | **-9.1** | **24.8** | **20.1** | **3.1** | **5.6** | **-5.0** |
| **Altai** | **-15.9** | **-17.5** | **-14.4** | **-9.2** | **12.7** | **13.1** | -1.1 |  | **3.2** | **-8.8** | **22.1** | **19.2** | 1.2 | **4.8** | **-5.2** |
| **Chukotka** | **-18.4** | **-21.6** | **-17.0** | **-12.0** | **8.4** | **10.3** | **-5.0** | **-3.2** |  | **-11.8** | **20.1** | **17.5** | **-2.7** | **2.5** | **-6.5** |
| **Bryansk** | **-7.5** | **-9.4** | **-7.1** | -1.1 | **20.3** | **17.7** | **9.1** | **8.8** | **11.8** |  | **26.6** | **23.1** | **11.1** | **9.5** | -1.0 |
| **Shanxi** | **-30.1** | **-33.4** | **-28.5** | **-27.3** | **-14.9** | **-10.8** | **-24.8** | **-22.1** | **-20.1** | **-26.6** |  | 1.7 | **-23.0** | **-14.0** | **-18.8** |
| **China** | **-26.2** | **-26.7** | **-25.2** | **-23.0** | **-13.5** | **-11.2** | **-20.1** | **-19.2** | **-17.5** | **-23.1** | -1.7 |  | **-18.9** | **-13.7** | **-19.7** |
| **Xinjiang_Marsden** | **-18.2** | **-21.8** | **-16.4** | **-11.8** | **12.0** | **13.2** | **-3.1** | -1.2 | **2.7** | **-11.1** | **23.0** | **18.9** |  | **4.4** | **-5.8** |
| **Qinghai** | **-14.2** | **-15.1** | **-14.8** | **-10.1** | **3.2** | **5.4** | **-5.6** | **-4.8** | **-2.5** | **-9.5** | **14.0** | **13.7** | **-4.4** |  | **-11.8** |
| **Tibet** | -2.4 | **-2.7** | -2.5 | 0.5 | **11.6** | **13.1** | **5.0** | **5.2** | **6.5** | 1.0 | **18.8** | **19.7** | **5.8** | **11.8** |  |
| **D(O Guizhou2;P3 P4)** | | | |  |  |  |  |  |  |  |  |  |  |  |  |
| **P3/P4** | **Taimyr** | **Iranwolf** | **Indianwolf** | **Portugal** | **Inner_Mongolia** | **Liaoning** | **Xinjiang** | **Altai** | **Chukotka** | **Bryansk** | **Shanxi** | **China** | **Xinjiang_Marsden** | **Qinghai** | **Tibet** |
| **Taimyr** |  | -0.1 | -0.6 | **6.5** | **26.5** | **26.5** | **18.8** | **15.8** | **18.6** | **8.8** | **30.4** | **28.3** | **19.1** | **13.1** | **3.4** |
| **Iranwolf** | 0.1 |  | -0.7 | **8.4** | **28.1** | **26.7** | **25.7** | **18.6** | **20.9** | **9.9** | **31.8** | **27.4** | **23.2** | **13.9** | **3.5** |
| **Indianwolf** | 0.6 | 0.7 |  | **7.2** | **24.7** | **23.0** | **18.7** | **16.0** | **18.5** | **8.4** | **29.4** | **26.2** | **18.9** | **14.1** | **4.0** |
| **Portugal** | **-6.5** | **-8.4** | **-7.2** |  | **19.0** | **19.9** | **11.0** | **9.5** | **11.9** | 1.3 | **25.4** | **23.6** | **12.3** | **9.3** | 0.3 |
| **Inner_Mongolia** | **-26.5** | **-28.1** | **-24.7** | **-19.0** |  | **5.5** | **-14.0** | **-11.3** | **-7.1** | **-19.8** | **15.2** | **13.5** | **-10.3** | **-2.7** | **-10.2** |
| **Liaoning** | **-26.5** | **-26.7** | **-23.0** | **-19.9** | **-5.5** |  | **-15.2** | **-13.5** | **-11.3** | **-20.0** | **9.1** | **9.2** | **-13.3** | **-6.1** | **-11.9** |
| **Xinjiang** | **-18.8** | **-25.7** | **-18.7** | **-11.0** | **14.0** | **15.2** |  | 0.0 | **4.4** | **-10.7** | **22.5** | **19.6** | **2.9** | **4.5** | **-4.4** |
| **Altai** | **-15.8** | **-18.6** | **-16.0** | **-9.5** | **11.3** | **13.5** | 0.0 |  | **3.6** | **-8.7** | **21.1** | **18.8** | 2.1 | **4.2** | **-4.2** |
| **Chukotka** | **-18.6** | **-20.9** | **-18.5** | **-11.9** | **7.1** | **11.3** | **-4.4** | **-3.6** |  | **-11.7** | **19.2** | **17.5** | -2.1 | 1.9 | **-5.9** |
| **Bryansk** | **-8.8** | **-9.9** | **-8.4** | -1.3 | **19.8** | **20.0** | **10.7** | **8.7** | **11.7** |  | **25.2** | **23.6** | **12.4** | **9.0** | -0.2 |
| **Shanxi** | **-30.4** | **-31.8** | **-29.4** | **-25.4** | **-15.2** | **-9.1** | **-22.5** | **-21.1** | **-19.2** | **-25.2** |  | 0.5 | **-20.5** | **-13.9** | **-18.2** |
| **China** | **-28.3** | **-27.4** | **-26.2** | **-23.6** | **-13.5** | **-9.2** | **-19.6** | **-18.8** | **-17.5** | **-23.6** | -0.5 |  | **-17.9** | **-12.9** | **-17.5** |
| **Xinjiang_Marsden** | **-19.1** | **-23.2** | **-18.9** | **-12.3** | **10.3** | **13.3** | **-2.9** | -2.1 | 2.1 | **-12.4** | **20.5** | **17.9** |  | **3.2** | **-5.0** |
| **Qinghai** | **-13.1** | **-13.9** | **-14.1** | **-9.3** | **2.7** | **6.1** | **-4.5** | **-4.2** | -1.9 | **-9.0** | **13.9** | **12.9** | **-3.2** |  | **-9.6** |
| **Tibet** | **-3.4** | **-3.5** | **-4.0** | -0.3 | **10.2** | **11.9** | **4.4** | **4.2** | **5.9** | 0.2 | **18.2** | **17.5** | **5.0** | **9.6** |  |
| **D(O Jiangxi;P3 P4)** | | | |  |  |  |  |  |  |  |  |  |  |  |  |
| **P3/P4** | **Taimyr** | **Iranwolf** | **Indianwolf** | **Portugal** | **Inner_Mongolia** | **Liaoning** | **Xinjiang** | **Altai** | **Chukotka** | **Bryansk** | **Shanxi** | **China** | **Xinjiang_Marsden** | **Qinghai** | **Tibet** |
| **Taimyr** |  | 0.7 | 1.4 | **3.0** | **22.6** | **24.2** | **17.5** | **14.7** | **15.9** | **5.1** | **27.1** | **23.5** | **16.7** | **24.0** | **24.1** |
| **Iranwolf** | -0.7 |  | 1.1 | **2.9** | **22.0** | **24.9** | **24.5** | **14.4** | **17.4** | **4.9** | **27.7** | **22.6** | **19.3** | **24.9** | **24.7** |
| **Indianwolf** | -1.4 | -1.1 |  | 1.6 | **18.5** | **20.5** | **15.9** | **11.7** | **13.4** | **3.0** | **23.7** | **20.3** | **13.7** | **24.6** | **24.8** |
| **Portugal** | **-3.0** | **-2.9** | -1.6 |  | **17.1** | **19.0** | **13.8** | **10.2** | **12.1** | 1.5 | **23.4** | **20.1** | **13.0** | **22.7** | **23.1** |
| **Inner_Mongolia** | **-22.6** | **-22.0** | **-18.5** | **-17.1** |  | **4.2** | **-9.0** | **-7.5** | **-6.6** | **-19.4** | **11.0** | **9.5** | **-9.0** | **17.4** | **19.6** |
| **Liaoning** | **-24.2** | **-24.9** | **-20.5** | **-19.0** | **-4.2** |  | **-13.0** | **-10.7** | **-10.6** | **-20.1** | **6.0** | **6.7** | **-12.0** | **15.9** | **18.6** |
| **Xinjiang** | **-17.5** | **-24.5** | **-15.9** | **-13.8** | **9.0** | **13.0** |  | -0.2 | 1.3 | **-14.1** | **18.1** | **15.2** | -0.1 | **20.9** | **21.8** |
| **Altai** | **-14.7** | **-14.4** | **-11.7** | **-10.2** | **7.5** | **10.7** | 0.2 |  | 1.2 | **-10.4** | **15.2** | **14.0** | 0.1 | **20.0** | **21.3** |
| **Chukotka** | **-15.9** | **-17.4** | **-13.4** | **-12.1** | **6.6** | **10.6** | -1.3 | -1.2 |  | **-12.2** | **16.1** | **13.9** | -1.3 | **19.2** | **20.5** |
| **Bryansk** | **-5.1** | **-4.9** | **-3.0** | -1.5 | **19.4** | **20.1** | **14.1** | **10.4** | **12.2** |  | **24.9** | **21.3** | **13.0** | **23.1** | **23.3** |
| **Shanxi** | **-27.1** | **-27.7** | **-23.7** | **-23.4** | **-11.0** | **-6.0** | **-18.1** | **-15.2** | **-16.1** | **-24.9** |  | 1.7 | **-17.0** | **12.9** | **16.3** |
| **China** | **-23.5** | **-22.6** | **-20.3** | **-20.1** | **-9.5** | **-6.7** | **-15.2** | **-14.0** | **-13.9** | **-21.3** | -1.7 |  | **-14.6** | **11.7** | **15.4** |
| **Xinjiang_Marsden** | **-16.7** | **-19.3** | **-13.7** | **-13.0** | **9.0** | **12.0** | 0.1 | -0.1 | 1.3 | **-13.0** | **17.0** | **14.6** |  | **20.2** | **21.3** |
| **Qinghai** | **-24.0** | **-24.9** | **-24.6** | **-22.7** | **-17.4** | **-15.9** | **-20.9** | **-20.0** | **-19.2** | **-23.1** | **-12.9** | **-11.7** | **-20.2** |  | **11.7** |
| **Tibet** | **-24.1** | **-24.7** | **-24.8** | **-23.1** | **-19.6** | **-18.6** | **-21.8** | **-21.3** | **-20.5** | **-23.3** | **-16.3** | **-15.4** | **-21.3** | **-11.7** |  |
| **D(O Zhejiang;P3 P4)** | | | |  |  |  |  |  |  |  |  |  |  |  |  |
| **P3/P4** | **Taimyr** | **Iranwolf** | **Indianwolf** | **Portugal** | **Inner_Mongolia** | **Liaoning** | **Xinjiang** | **Altai** | **Chukotka** | **Bryansk** | **Shanxi** | **China** | **Xinjiang_Marsden** | **Qinghai** | **Tibet** |
| **Taimyr** |  | -0.7 | -0.6 | **5.3** | **21.3** | **23.1** | **13.7** | **11.7** | **14.5** | **4.6** | **31.4** | **28.0** | **13.8** | **16.5** | **7.4** |
| **Iranwolf** | 0.7 |  | -0.1 | **8.7** | **27.0** | **27.8** | **21.4** | **15.3** | **20.5** | **7.7** | **38.0** | **29.4** | **18.5** | **19.4** | **8.4** |
| **Indianwolf** | 0.6 | 0.1 |  | **7.1** | **21.5** | **22.6** | **15.2** | **12.6** | **15.6** | **5.9** | **30.9** | **27.3** | **15.0** | **18.7** | **8.5** |
| **Portugal** | **-5.3** | **-8.7** | **-7.1** |  | **15.4** | **17.5** | **7.5** | **6.3** | **9.8** | -0.9 | **28.3** | **23.5** | **8.2** | **13.2** | **4.5** |
| **Inner_Mongolia** | **-21.3** | **-27.0** | **-21.5** | **-15.4** |  | **4.7** | **-13.8** | **-11.7** | **-6.9** | **-18.7** | **16.4** | **14.6** | **-10.4** | 1.8 | **-4.4** |
| **Liaoning** | **-23.1** | **-27.8** | **-22.6** | **-17.5** | **-4.7** |  | **-16.6** | **-14.3** | **-10.5** | **-19.9** | **11.0** | **12.4** | **-14.2** | -1.6 | **-6.7** |
| **Xinjiang** | **-13.7** | **-21.4** | **-15.2** | **-7.5** | **13.8** | **16.6** |  | 0.1 | **4.8** | **-9.7** | **28.3** | **22.7** | 2.1 | **9.9** | 1.3 |
| **Altai** | **-11.7** | **-15.3** | **-12.6** | **-6.3** | **11.7** | **14.3** | -0.1 |  | **3.7** | **-8.3** | **25.2** | **21.5** | 1.4 | **9.1** | 1.2 |
| **Chukotka** | **-14.5** | **-20.5** | **-15.6** | **-9.8** | **6.9** | **10.5** | **-4.8** | **-3.7** |  | **-11.3** | **20.7** | **18.7** | **-2.9** | **6.1** | -0.7 |
| **Bryansk** | **-4.6** | **-7.7** | **-5.9** | 0.9 | **18.7** | **19.9** | **9.7** | **8.3** | **11.3** |  | **30.0** | **25.3** | **10.0** | **13.9** | **4.9** |
| **Shanxi** | **-31.4** | **-38.0** | **-30.9** | **-28.3** | **-16.4** | **-11.0** | **-28.3** | **-25.2** | **-20.7** | **-30.0** |  | **2.6** | **-25.0** | **-9.7** | **-12.5** |
| **China** | **-28.0** | **-29.4** | **-27.3** | **-23.5** | **-14.6** | **-12.4** | **-22.7** | **-21.5** | **-18.7** | **-25.3** | **-2.6** |  | **-21.5** | **-10.5** | **-14.2** |
| **Xinjiang_Marsden** | **-13.8** | **-18.5** | **-15.0** | **-8.2** | **10.4** | **14.2** | -2.1 | -1.4 | **2.9** | **-10.0** | **25.0** | **21.5** |  | **8.7** | 0.7 |
| **Qinghai** | **-16.5** | **-19.4** | **-18.7** | **-13.2** | -1.8 | 1.6 | **-9.9** | **-9.1** | **-6.1** | **-13.9** | **9.7** | **10.5** | **-8.7** |  | **-8.2** |
| **Tibet** | **-7.4** | **-8.4** | **-8.5** | **-4.5** | **4.4** | **6.7** | -1.3 | -1.2 | 0.7 | **-4.9** | **12.5** | **14.2** | -0.7 | **8.2** |  |
| **D(O Jilin;P3 P4)** | | |  |  |  |  |  |  |  |  |  |  |  |  |  |
| **P3/P4** | **Taimyr** | **Iranwolf** | **Indianwolf** | **Portugal** | **Inner_Mongolia** | **Liaoning** | **Xinjiang** | **Altai** | **Chukotka** | **Bryansk** | **Shanxi** | **China** | **Xinjiang_Marsden** | **Qinghai** | **Tibet** |
| **Taimyr** |  | -1.2 | -2.0 | **5.4** | **25.3** | **21.8** | **14.3** | **15.0** | **18.0** | **7.6** | **23.6** | **21.2** | **16.3** | **6.8** | -1.2 |
| **Iranwolf** | 1.2 |  | -1.4 | **8.6** | **29.1** | **23.7** | **24.3** | **15.6** | **22.4** | **10.2** | **26.5** | **21.4** | **20.7** | **8.0** | -0.6 |
| **Indianwolf** | 2.0 | 1.4 |  | **7.6** | **23.3** | **21.3** | **17.9** | **13.7** | **19.2** | **8.7** | **22.0** | **20.2** | **16.6** | **8.5** | -0.1 |
| **Portugal** | **-5.4** | **-8.6** | **-7.6** |  | **18.4** | **16.9** | **9.7** | **8.2** | **14.1** | 1.8 | **18.0** | **15.4** | **10.1** | **3.4** | **-4.0** |
| **Inner_Mongolia** | **-25.3** | **-29.1** | **-23.3** | **-18.4** |  | **3.9** | **-15.5** | **-10.0** | **-5.3** | **-18.8** | **3.0** | 2.4 | **-13.5** | **-8.8** | **-14.0** |
| **Liaoning** | **-21.8** | **-23.7** | **-21.3** | **-16.9** | **-3.9** |  | **-14.2** | **-11.1** | **-7.2** | **-16.7** | -1.2 | -1.1 | **-13.1** | **-10.8** | **-15.6** |
| **Xinjiang** | **-14.3** | **-24.3** | **-17.9** | **-9.7** | **15.5** | **14.2** |  | 2.3 | **7.5** | **-7.1** | **15.3** | **12.2** | 1.8 | -1.5 | **-8.3** |
| **Altai** | **-15.0** | **-15.6** | **-13.7** | **-8.2** | **10.0** | **11.1** | -2.3 |  | **3.7** | **-8.0** | **11.7** | **9.2** | -1.1 | -2.5 | **-8.7** |
| **Chukotka** | **-18.0** | **-22.4** | **-19.2** | **-14.1** | **5.3** | **7.2** | **-7.5** | **-3.7** |  | **-12.0** | **6.9** | **5.9** | **-5.7** | **-5.2** | **-11.0** |
| **Bryansk** | **-7.6** | **-10.2** | **-8.7** | -1.8 | **18.8** | **16.7** | **7.1** | **8.0** | **12.0** |  | **18.8** | **15.6** | **8.1** | 2.1 | **-4.8** |
| **Shanxi** | **-23.6** | **-26.5** | **-22.0** | **-18.0** | **-3.0** | 1.2 | **-15.3** | **-11.7** | **-6.9** | **-18.8** |  | 0.0 | **-13.6** | **-10.4** | **-14.8** |
| **China** | **-21.2** | **-21.4** | **-20.2** | **-15.4** | -2.4 | 1.1 | **-12.2** | **-9.2** | **-5.9** | **-15.6** | 0.0 |  | **-11.1** | **-10.1** | **-14.7** |
| **Xinjiang_Marsden** | **-16.3** | **-20.7** | **-16.6** | **-10.1** | **13.5** | **13.1** | -1.8 | 1.1 | **5.7** | **-8.1** | **13.6** | **11.1** |  | -2.1 | **-8.8** |
| **Qinghai** | **-6.8** | **-8.0** | **-8.5** | **-3.4** | **8.8** | **10.8** | 1.5 | 2.5 | **5.2** | -2.1 | **10.4** | **10.1** | 2.1 |  | **-9.8** |
| **Tibet** | 1.2 | 0.6 | 0.1 | **4.0** | **14.0** | **15.6** | **8.3** | **8.7** | **11.0** | **4.8** | **14.8** | **14.7** | **8.8** | **9.8** |  |
| **D(O Heilongjiang;P3 P4)** | | | |  |  |  |  |  |  |  |  |  |  |  |  |
| **P3/P4** | **Taimyr** | **Iranwolf** | **Indianwolf** | **Portugal** | **Inner_Mongolia** | **Liaoning** | **Xinjiang** | **Altai** | **Chukotka** | **Bryansk** | **Shanxi** | **China** | **Xinjiang_Marsden** | **Qinghai** | **Tibet** |
| **Taimyr** |  | -1.0 | -1.1 | **5.5** | **22.9** | **23.9** | **13.9** | **13.1** | **16.0** | **7.7** | **23.3** | **20.7** | **15.7** | **5.9** | -2.4 |
| **Iranwolf** | 1.0 |  | -0.5 | **8.5** | **25.4** | **25.3** | **24.4** | **14.8** | **20.4** | **9.3** | **24.3** | **21.8** | **20.8** | **7.4** | -1.9 |
| **Indianwolf** | 1.1 | 0.5 |  | **6.6** | **21.1** | **22.1** | **16.2** | **12.4** | **16.2** | **7.2** | **21.0** | **19.4** | **15.4** | **7.3** | -1.7 |
| **Portugal** | **-5.5** | **-8.5** | **-6.6** |  | **15.6** | **18.9** | **8.7** | **7.4** | **12.2** | 1.6 | **16.0** | **14.6** | **10.1** | **2.6** | **-5.1** |
| **Inner_Mongolia** | **-22.9** | **-25.4** | **-21.1** | **-15.6** |  | **7.0** | **-12.8** | **-9.4** | **-5.3** | **-16.4** | 2.3 | **3.0** | **-10.0** | **-9.7** | **-14.7** |
| **Liaoning** | **-23.9** | **-25.3** | **-22.1** | **-18.9** | **-7.0** |  | **-15.9** | **-13.9** | **-10.3** | **-19.3** | **-5.0** | **-3.8** | **-14.6** | **-12.9** | **-16.6** |
| **Xinjiang** | **-13.9** | **-24.4** | **-16.2** | **-8.7** | **12.8** | **15.9** |  | 1.3 | **6.4** | **-6.1** | **13.2** | **12.0** | **2.7** | **-2.6** | **-9.3** |
| **Altai** | **-13.1** | **-14.8** | **-12.4** | **-7.4** | **9.4** | **13.9** | -1.3 |  | **4.0** | **-7.4** | **11.4** | **10.2** | 0.4 | **-2.9** | **-9.1** |
| **Chukotka** | **-16.0** | **-20.4** | **-16.2** | **-12.2** | **5.3** | **10.3** | **-6.4** | **-4.0** |  | **-10.1** | **7.0** | **6.9** | **-4.4** | **-5.8** | **-11.3** |
| **Bryansk** | **-7.7** | **-9.3** | **-7.2** | -1.6 | **16.4** | **19.3** | **6.1** | **7.4** | **10.1** |  | **17.0** | **15.6** | **7.8** | 1.3 | **-5.7** |
| **Shanxi** | **-23.3** | **-24.3** | **-21.0** | **-16.0** | -2.3 | **5.0** | **-13.2** | **-11.4** | **-7.0** | **-17.0** |  | 1.0 | **-10.8** | **-10.6** | **-15.3** |
| **China** | **-20.7** | **-21.8** | **-19.4** | **-14.6** | **-3.0** | **3.8** | **-12.0** | **-10.2** | **-6.9** | **-15.6** | -1.0 |  | **-10.1** | **-10.5** | **-15.0** |
| **Xinjiang_Marsden** | **-15.7** | **-20.8** | **-15.4** | **-10.1** | **10.0** | **14.6** | **-2.7** | -0.4 | **4.4** | **-7.8** | **10.8** | **10.1** |  | **-3.6** | **-10.1** |
| **Qinghai** | **-5.9** | **-7.4** | **-7.3** | **-2.6** | **9.7** | **12.9** | **2.6** | **2.9** | **5.8** | -1.3 | **10.6** | **10.5** | **3.6** |  | **-10.2** |
| **Tibet** | 2.4 | 1.9 | 1.7 | **5.1** | **14.7** | **16.6** | **9.3** | **9.1** | **11.3** | **5.7** | **15.3** | **15.0** | **10.1** | **10.2** |  |

Table S4. F3 tests among the clades among SEA, NEA, and Tibetan gray wolves.

| **Source 1** | **Source 1** | **Target** | **f_3** | **std. err** | **Z** | **SNPs** |
| --- | --- | --- | --- | --- | --- | --- |
| Tibetan | SEA | Jiangxi | -0.08023 | 0.007068 | -11.352 | 2638661 |
| Tibetan | NEA | Jiangxi | -0.06734 | 0.007366 | -9.141 | 2478583 |
| Tibetan | SEA | Qinghai | -0.09735 | 0.005096 | -19.101 | 2631539 |
| Tibetan | NEA | Qinghai | -0.09298 | 0.00522 | -17.811 | 2488007 |

Table S5. Z-score of *D(Fox, X; test, Y).*

| **D(O P2;Guizhou1 P4)** | | | |  |  |  |  |  |  |  |  |  |  |  |  |
| --- | --- | --- | --- | --- | --- | --- | --- | --- | --- | --- | --- | --- | --- | --- | --- |
| **P2/P4** | **Taimyr** | **Iranwolf** | **Indianwolf** | **Portugal** | **Inner_Mongolia** | **Liaoning** | **Xinjiang** | **Altai** | **Chukotka** | **Bryansk** | **Shanxi** | **China** | **Xinjiang_Marsden** | **Qinghai** | **Tibet** |
| **Taimyr** |  | **-5.5** | **-6.7** | **3.8** | **2.7** | 2.0 | -0.3 | 2.4 | **4.1** | **2.6** | 2.1 | 1.8 | 1.1 | **-8.5** | **-12.5** |
| **Iranwolf** | **-4.4** |  | **25.0** | **15.1** | **4.6** | **2.8** | **14.1** | **11.7** | **8.2** | **15.4** | 2.1 | 2.0 | **13.4** | **-6.3** | **-10.9** |
| **Indianwolf** | **-5.2** | **24.5** |  | **9.0** | **3.6** | 1.5 | **11.4** | **8.9** | **6.2** | **8.8** | 0.4 | -0.2 | **9.9** | **-4.5** | **-8.6** |
| **Portugal** | -1.8 | **8.8** | **4.4** |  | **4.6** | **2.9** | **11.7** | **8.2** | **8.3** | **19.3** | 1.0 | -0.6 | **10.3** | **-7.3** | **-14.6** |
| **Inner_Mongolia** | **-22.1** | **-26.4** | **-21.7** | **-14.2** |  | 2.2 | **-10.3** | **-7.0** | **-3.4** | **-13.2** | 2.2 | **2.9** | **-7.4** | **-10.3** | **-14.3** |
| **Liaoning** | **-21.4** | **-23.8** | **-20.4** | **-16.6** | -1.7 |  | **-13.2** | **-10.5** | **-7.0** | **-14.8** | 1.3 | 1.8 | **-10.8** | **-10.8** | **-14.9** |
| **Xinjiang** | **-17.2** | **-5.7** | **-4.7** | 2.1 | **7.4** | **3.7** |  | **7.3** | **4.6** | 1.2 | **3.1** | **3.5** | **17.6** | **-6.8** | **-11.0** |
| **Altai** | **-11.6** | **-4.7** | **-4.6** | 0.7 | **5.0** | 2.0 | **5.4** |  | **5.0** | **3.1** | 2.3 | **3.5** | **7.0** | **-6.7** | **-11.5** |
| **Chukotka** | **-12.3** | **-11.1** | **-10.1** | -2.2 | **5.2** | **4.6** | -0.4 | 1.8 |  | -2.5 | 1.7 | **2.8** | 1.0 | **-8.4** | **-12.9** |
| **Bryansk** | **-4.2** | **7.8** | **3.4** | **19.7** | **4.8** | **3.4** | **9.4** | **10.4** | **7.6** |  | 1.9 | 1.8 | **10.4** | **-7.8** | **-12.5** |
| **Shanxi** | **-28.4** | **-31.0** | **-28.3** | **-26.1** | **-13.3** | **-9.4** | **-22.5** | **-20.2** | **-19.3** | **-24.2** |  | **4.1** | **-21.5** | **-13.8** | **-17.9** |
| **China** | **-25.4** | **-26.7** | **-26.3** | **-23.7** | **-12.0** | **-9.7** | **-19.5** | **-17.4** | **-16.4** | **-22.0** | 2.3 |  | **-19.6** | **-13.4** | **-19.0** |
| **Xinjiang_Marsden** | **-16.8** | **-5.8** | **-7.3** | -0.6 | **5.9** | **3.0** | **14.9** | **6.9** | **3.9** | 1.2 | 2.0 | 0.7 |  | **-9.2** | **-12.6** |
| **Qinghai** | **-23.2** | **-22.7** | **-18.5** | **-18.4** | **-6.1** | **-4.7** | **-13.7** | **-11.4** | **-11.8** | **-18.7** | 0.3 | -0.8 | **-14.4** |  | **22.3** |
| **Tibet** | **-18.9** | **-18.3** | **-14.4** | **-17.4** | -2.4 | -1.8 | **-7.3** | **-7.9** | **-8.2** | **-16.8** | 2.0 | 0.7 | **-9.4** | **29.4** |  |
| **D(O P2;Guizhou2 P4)** | | | |  |  |  |  |  |  |  |  |  |  |  |  |
| **P2/P4** | **Taimyr** | **Iranwolf** | **Indianwolf** | **Portugal** | **Inner_Mongolia** | **Liaoning** | **Xinjiang** | **Altai** | **Chukotka** | **Bryansk** | **Shanxi** | **China** | **Xinjiang_Marsden** | **Qinghai** | **Tibet** |
| **Taimyr** |  | **-4.9** | **-5.8** | **3.6** | **2.7** | 2.1 | 0.0 | 2.3 | **3.5** | 2.5 | 1.9 | 1.8 | 1.0 | **-7.7** | **-12.1** |
| **Iranwolf** | **-4.8** |  | **22.6** | **14.5** | **4.0** | 2.4 | **13.5** | **11.0** | **7.4** | **14.2** | 1.7 | 1.4 | **13.2** | **-6.7** | **-11.5** |
| **Indianwolf** | **-4.9** | **23.2** |  | **9.8** | **4.1** | 2.0 | **12.4** | **9.2** | **6.7** | **8.9** | 1.0 | 0.4 | **11.0** | **-4.1** | **-8.7** |
| **Portugal** | -1.8 | **8.6** | **4.2** |  | **4.3** | **2.8** | **11.6** | **8.0** | **8.1** | **19.9** | 0.8 | -0.7 | **9.5** | **-7.9** | **-15.2** |
| **Inner_Mongolia** | **-21.7** | **-24.6** | **-18.7** | **-12.8** |  | **3.5** | **-8.3** | **-6.0** | -2.2 | **-12.6** | **3.6** | **4.2** | **-5.8** | **-9.6** | **-14.0** |
| **Liaoning** | **-21.9** | **-23.9** | **-20.4** | **-17.3** | -2.4 |  | **-12.9** | **-11.3** | **-7.9** | **-15.7** | 0.5 | 1.3 | **-11.5** | **-10.7** | **-14.7** |
| **Xinjiang** | **-16.3** | **-6.3** | **-5.1** | 1.2 | **5.6** | **2.5** |  | **6.3** | **3.6** | 0.3 | 1.8 | 2.3 | **15.4** | **-7.6** | **-11.7** |
| **Altai** | **-10.7** | **-4.7** | **-4.2** | 0.8 | **5.0** | 2.2 | **5.4** |  | **5.1** | **3.2** | 2.4 | **3.4** | **7.1** | **-6.6** | **-11.5** |
| **Chukotka** | **-11.7** | **-11.2** | **-9.7** | **-2.5** | **4.6** | **4.0** | -0.8 | 1.5 |  | **-2.8** | 1.3 | 2.4 | 0.7 | **-8.7** | **-13.8** |
| **Bryansk** | **-4.2** | **7.2** | **2.9** | **20.6** | **4.9** | **3.2** | **9.0** | **10.5** | **7.3** |  | 1.7 | 1.4 | **10.2** | **-8.9** | **-13.9** |
| **Shanxi** | **-27.2** | **-29.3** | **-27.1** | **-24.1** | **-11.8** | **-8.5** | **-20.6** | **-18.8** | **-18.9** | **-22.9** |  | **4.2** | **-19.4** | **-13.2** | **-17.8** |
| **China** | **-26.3** | **-26.4** | **-25.2** | **-23.7** | **-10.6** | **-7.9** | **-18.8** | **-16.0** | **-15.7** | **-21.6** | **3.7** |  | **-18.6** | **-12.6** | **-17.6** |
| **Xinjiang_Marsden** | **-15.6** | **-6.4** | **-7.6** | -1.2 | **4.3** | 1.9 | **13.2** | **6.0** | **3.1** | 0.5 | 0.9 | 0.0 |  | **-9.6** | **-13.0** |
| **Qinghai** | **-21.7** | **-21.0** | **-16.3** | **-16.7** | **-5.3** | **-4.2** | **-12.1** | **-10.4** | **-10.6** | **-17.8** | 0.7 | -0.3 | **-13.0** |  | **22.0** |
| **Tibet** | **-19.4** | **-18.2** | **-14.4** | **-17.9** | **-3.8** | **-3.0** | **-7.9** | **-8.8** | **-9.1** | **-17.1** | 0.5 | -0.7 | **-9.6** | **28.6** |  |
| **D(O P2;Jiangxi P4)** | | | |  |  |  |  |  |  |  |  |  |  |  |  |
| **P2/P4** | **Taimyr** | **Iranwolf** | **Indianwolf** | **Portugal** | **Inner_Mongolia** | **Liaoning** | **Xinjiang** | **Altai** | **Chukotka** | **Bryansk** | **Shanxi** | **China** | **Xinjiang_Marsden** | **Qinghai** | **Tibet** |
| **Taimyr** |  | **3.6** | 2.2 | **8.4** | **8.6** | **8.6** | **7.3** | **8.1** | **8.2** | **7.9** | **8.2** | **8.3** | **7.5** | -0.5 | **-8.3** |
| **Iranwolf** | **3.2** |  | **22.5** | **15.9** | **9.2** | **8.4** | **14.9** | **12.9** | **10.8** | **15.9** | **7.5** | **7.6** | **14.2** | 0.9 | **-6.9** |
| **Indianwolf** | 1.2 | **21.9** |  | **12.3** | **8.1** | **6.9** | **14.1** | **11.7** | **9.9** | **11.4** | **6.1** | **5.5** | **12.6** | 1.6 | **-4.8** |
| **Portugal** | **6.0** | **14.0** | **11.0** |  | **10.9** | **10.1** | **17.1** | **13.2** | **14.4** | **22.3** | **8.6** | **7.9** | **14.8** | 0.9 | **-8.7** |
| **Inner_Mongolia** | **-6.3** | **-5.8** | **-5.1** | -0.8 |  | **10.1** | **4.6** | **4.5** | **7.2** | -0.3 | **10.2** | **11.0** | **5.3** | 1.8 | **-7.9** |
| **Liaoning** | **-9.0** | **-9.2** | **-9.3** | **-4.5** | **7.3** |  | 0.1 | 0.3 | **3.8** | **-3.7** | **9.1** | **9.7** | 1.1 | -0.6 | **-9.2** |
| **Xinjiang** | **-3.5** | **3.9** | **4.0** | **8.4** | **11.2** | **9.5** |  | **11.4** | **9.8** | **7.7** | **9.1** | **9.8** | **18.5** | 1.8 | **-6.3** |
| **Altai** | -1.3 | **3.3** | **3.3** | **6.6** | **9.6** | **7.9** | **10.5** |  | **9.5** | **8.1** | **8.4** | **9.2** | **11.3** | 1.0 | **-6.8** |
| **Chukotka** | -1.2 | 0.5 | 0.3 | **5.8** | **11.8** | **11.8** | **8.5** | **8.9** |  | **5.6** | **9.6** | **10.0** | **9.2** | 1.1 | **-7.4** |
| **Bryansk** | **5.0** | **13.0** | **9.7** | **22.3** | **11.1** | **10.7** | **14.9** | **15.0** | **12.9** |  | **9.9** | **9.7** | **14.7** | 0.9 | **-7.8** |
| **Shanxi** | **-12.1** | **-12.4** | **-12.9** | **-8.9** | **3.0** | **4.9** | **-4.4** | **-3.5** | -2.1 | **-8.1** |  | **13.8** | **-3.6** | -0.5 | **-10.2** |
| **China** | **-11.9** | **-12.8** | **-13.1** | **-10.4** | 2.0 | **3.9** | **-4.9** | **-3.6** | -2.4 | **-8.6** | **12.8** |  | **-5.0** | -2.3 | **-11.2** |
| **Xinjiang_Marsden** | -2.5 | **3.9** | **3.0** | **6.9** | **11.0** | **9.8** | **17.7** | **11.5** | **10.2** | **8.0** | **9.4** | **8.9** |  | 0.2 | **-7.6** |
| **Qinghai** | **-25.5** | **-25.8** | **-24.1** | **-23.0** | **-17.4** | **-17.2** | **-21.4** | **-20.6** | **-20.3** | **-23.6** | **-14.7** | **-14.7** | **-21.7** |  | **15.1** |
| **Tibet** | **-28.7** | **-28.7** | **-27.5** | **-28.3** | **-23.8** | **-23.9** | **-25.6** | **-25.5** | **-25.4** | **-27.7** | **-22.4** | **-23.2** | **-25.9** | **3.4** |  |
| **D(O P2;Zhejiang P4)** | | | |  |  |  |  |  |  |  |  |  |  |  |  |
| **P2/P4** | **Taimyr** | **Iranwolf** | **Indianwolf** | **Portugal** | **Inner_Mongolia** | **Liaoning** | **Xinjiang** | **Altai** | **Chukotka** | **Bryansk** | **Shanxi** | **China** | **Xinjiang_Marsden** | **Qinghai** | **Tibet** |
| **Taimyr** |  | **47.9** | **40.2** | **46.5** | **52.8** | **50.0** | **56.7** | **48.9** | **49.5** | **46.4** | **51.9** | **46.9** | **54.5** | **31.5** | **18.3** |
| **Iranwolf** | **43.4** |  | **65.2** | **60.1** | **61.0** | **54.5** | **74.4** | **59.8** | **59.9** | **57.8** | **58.1** | **52.5** | **70.1** | **35.5** | **21.9** |
| **Indianwolf** | **35.6** | **66.8** |  | **48.4** | **47.8** | **47.9** | **68.7** | **50.7** | **54.8** | **46.4** | **48.7** | **44.6** | **64.5** | **33.3** | **22.6** |
| **Portugal** | **37.4** | **55.2** | **43.2** |  | **46.3** | **46.6** | **66.7** | **47.2** | **53.9** | **50.1** | **47.3** | **42.9** | **62.3** | **28.1** | **17.3** |
| **Inner_Mongolia** | **28.5** | **38.1** | **31.5** | **37.8** |  | **53.1** | **58.7** | **46.8** | **50.4** | **37.4** | **57.0** | **55.3** | **53.9** | **35.7** | **18.4** |
| **Liaoning** | **23.1** | **27.6** | **22.8** | **29.8** | **42.8** |  | **41.3** | **34.3** | **42.2** | **28.7** | **46.9** | **43.9** | **40.4** | **29.8** | **15.0** |
| **Xinjiang** | **38.9** | **58.8** | **53.0** | **57.7** | **71.5** | **64.8** |  | **63.0** | **63.1** | **53.7** | **67.7** | **61.2** | **72.2** | **40.7** | **22.9** |
| **Altai** | **35.3** | **47.8** | **43.8** | **45.3** | **59.8** | **51.4** | **63.9** |  | **55.9** | **47.5** | **58.3** | **51.5** | **59.8** | **32.9** | **19.3** |
| **Chukotka** | **31.6** | **41.4** | **36.2** | **42.3** | **54.4** | **56.4** | **55.0** | **46.4** |  | **40.9** | **55.0** | **50.9** | **54.2** | **32.7** | **16.4** |
| **Bryansk** | **42.1** | **58.3** | **47.9** | **53.7** | **57.7** | **55.5** | **65.4** | **57.5** | **56.0** |  | **60.3** | **51.6** | **62.4** | **31.0** | **18.0** |
| **Shanxi** | **12.9** | **15.6** | **12.5** | **19.2** | **36.1** | **34.8** | **29.6** | **26.6** | **30.6** | **19.8** |  | **34.8** | **30.4** | **22.9** | **11.1** |
| **China** | **9.1** | **10.0** | **8.5** | **12.4** | **27.5** | **29.7** | **21.5** | **19.6** | **22.0** | **13.7** | **30.7** |  | **21.7** | **16.6** | **8.1** |
| **Xinjiang_Marsden** | **37.0** | **49.3** | **47.6** | **52.1** | **62.6** | **58.9** | **67.2** | **56.2** | **59.5** | **47.3** | **61.8** | **56.2** |  | **34.8** | **19.2** |
| **Qinghai** | **15.0** | **20.4** | **16.9** | **18.4** | **34.7** | **34.2** | **38.2** | **26.5** | **32.4** | **21.0** | **39.6** | **33.0** | **33.0** |  | **36.7** |
| **Tibet** | **14.4** | **20.2** | **19.2** | **17.0** | **32.8** | **32.4** | **34.8** | **25.7** | **26.0** | **20.6** | **34.4** | **32.3** | **29.9** | **41.5** |  |
| **D(O P2;Jilin P4)** | | |  |  |  |  |  |  |  |  |  |  |  |  |  |
| **P2/P4** | **Taimyr** | **Iranwolf** | **Indianwolf** | **Portugal** | **Inner_Mongolia** | **Liaoning** | **Xinjiang** | **Altai** | **Chukotka** | **Bryansk** | **Shanxi** | **China** | **Xinjiang_Marsden** | **Qinghai** | **Tibet** |
| **Taimyr** |  | **-6.0** | **-7.7** | **3.0** | 1.6 | 0.8 | -1.4 | 1.5 | **2.8** | 2.0 | 1.0 | 0.9 | -0.2 | **-8.5** | **-12.9** |
| **Iranwolf** | **-4.7** |  | **22.7** | **13.8** | **4.6** | 2.3 | **13.1** | **12.4** | **7.0** | **18.2** | 1.7 | 1.0 | **13.2** | **-5.9** | **-10.2** |
| **Indianwolf** | **-4.8** | **21.4** |  | **8.0** | **6.1** | **3.0** | **11.3** | **11.9** | **6.1** | **12.7** | 1.6 | 0.5 | **10.0** | **-3.0** | **-7.7** |
| **Portugal** | **-2.6** | **6.7** | **2.8** |  | **4.7** | **2.5** | **9.1** | **8.8** | **7.2** | **22.6** | 0.2 | -2.0 | **8.1** | **-6.7** | **-14.1** |
| **Inner_Mongolia** | **-25.0** | **-25.0** | **-19.4** | **-13.3** |  | **2.9** | **-9.3** | **-7.8** | **-3.2** | **-14.4** | **3.1** | **3.6** | **-7.1** | **-8.9** | **-13.4** |
| **Liaoning** | **-22.0** | **-23.5** | **-20.4** | **-15.3** | -1.2 |  | **-12.8** | **-11.6** | **-6.3** | **-16.0** | 1.8 | 2.3 | **-10.5** | **-9.4** | **-14.6** |
| **Xinjiang** | **-16.9** | **-6.3** | **-5.3** | 1.8 | **6.5** | **3.3** |  | **6.9** | **3.9** | 1.0 | 2.4 | **2.7** | **18.1** | **-6.9** | **-10.9** |
| **Altai** | **-13.1** | **-5.7** | **-5.3** | -0.5 | **3.0** | 0.3 | **3.2** |  | **3.2** | 1.9 | 0.4 | 1.5 | **5.1** | **-7.4** | **-12.2** |
| **Chukotka** | **-14.6** | **-13.4** | **-12.4** | **-4.2** | 2.4 | 1.5 | **-3.4** | -0.7 |  | **-5.1** | -1.3 | -0.2 | -1.8 | **-10.5** | **-14.8** |
| **Bryansk** | **-5.9** | **6.0** | 1.6 | **18.4** | **3.8** | 1.6 | **7.7** | **10.3** | **6.4** |  | 0.1 | -0.4 | **9.0** | **-8.5** | **-13.5** |
| **Shanxi** | **-24.2** | **-25.8** | **-22.5** | **-17.9** | 0.0 | **3.3** | **-13.0** | **-11.9** | **-8.7** | **-18.9** |  | **15.7** | **-10.9** | **-4.0** | **-10.8** |
| **China** | **-21.7** | **-23.5** | **-21.6** | **-17.5** | 0.6 | **3.5** | **-12.1** | **-9.1** | **-6.5** | **-16.1** | **15.9** |  | **-11.4** | **-4.9** | **-11.4** |
| **Xinjiang_Marsden** | **-17.7** | **-6.2** | **-7.2** | -0.1 | **6.2** | **3.4** | **16.3** | **7.9** | **4.5** | 1.9 | **2.5** | 0.7 |  | **-8.8** | **-12.4** |
| **Qinghai** | **-20.5** | **-15.1** | **-10.7** | **-10.5** | 1.6 | 2.4 | **-5.5** | **-6.6** | **-4.8** | **-16.2** | **8.4** | **6.4** | **-6.8** |  | **24.6** |
| **Tibet** | **-17.8** | **-16.5** | **-11.3** | **-14.6** | 2.5 | 2.3 | **-2.8** | **-5.1** | **-5.2** | **-15.7** | **6.8** | **4.3** | **-5.5** | **30.8** |  |
| **D(O P2;Heilongjiang P4)** | | | |  |  |  |  |  |  |  |  |  |  |  |  |
| **P2/P4** | **Taimyr** | **Iranwolf** | **Indianwolf** | **Portugal** | **Inner_Mongolia** | **Liaoning** | **Xinjiang** | **Altai** | **Chukotka** | **Bryansk** | **Shanxi** | **China** | **Xinjiang_Marsden** | **Qinghai** | **Tibet** |
| **Taimyr** |  | **-4.8** | **-7.1** | **2.7** | 2.3 | 1.5 | -0.2 | 2.0 | **3.5** | **2.5** | 1.7 | 1.4 | 0.7 | **-7.8** | **-12.1** |
| **Iranwolf** | **-3.9** |  | **23.3** | **14.1** | **4.9** | **2.9** | **14.3** | **12.2** | **7.5** | **19.1** | 2.0 | 0.9 | **14.3** | **-5.6** | **-10.0** |
| **Indianwolf** | **-5.5** | **20.3** |  | **7.0** | **5.1** | 2.1 | **10.6** | **9.9** | **5.4** | **10.8** | 0.7 | -0.3 | **9.2** | **-3.3** | **-7.7** |
| **Portugal** | **-2.9** | **6.8** | **2.8** |  | **4.7** | 2.4 | **9.1** | **8.5** | **7.4** | **23.9** | -0.5 | **-2.5** | **8.9** | **-7.0** | **-14.3** |
| **Inner_Mongolia** | **-23.2** | **-21.7** | **-18.3** | **-11.4** |  | **4.0** | **-7.4** | **-6.6** | -1.9 | **-13.0** | **4.2** | **4.7** | **-5.1** | **-8.0** | **-12.6** |
| **Liaoning** | **-22.7** | **-23.4** | **-21.6** | **-17.1** | **-3.6** |  | **-13.8** | **-13.5** | **-8.3** | **-17.6** | -0.8 | -0.3 | **-12.3** | **-11.2** | **-15.0** |
| **Xinjiang** | **-16.4** | **-5.5** | **-4.8** | 1.7 | **6.5** | **3.5** |  | **7.0** | **4.3** | 1.0 | **2.6** | 2.4 | **19.4** | **-6.7** | **-10.5** |
| **Altai** | **-12.2** | **-4.8** | **-5.2** | -0.2 | **4.0** | 1.0 | **4.2** |  | **4.0** | 2.2 | 1.2 | 2.1 | **6.6** | **-6.6** | **-11.3** |
| **Chukotka** | **-12.7** | **-12.3** | **-11.2** | **-3.2** | **3.7** | **3.0** | -2.0 | 0.3 |  | **-4.2** | -0.2 | 1.1 | -0.5 | **-9.4** | **-14.0** |
| **Bryansk** | **-5.8** | **6.0** | 1.6 | **18.4** | **3.9** | 1.9 | **7.4** | **9.7** | **6.2** |  | -0.5 | -0.8 | **9.3** | **-9.1** | **-13.7** |
| **Shanxi** | **-24.1** | **-23.7** | **-22.4** | **-17.4** | 1.5 | **4.9** | **-11.6** | **-9.8** | **-7.6** | **-18.5** |  | **16.5** | **-9.7** | **-3.1** | **-10.0** |
| **China** | **-21.4** | **-23.7** | **-21.7** | **-18.0** | 1.4 | **4.1** | **-11.7** | **-9.3** | **-6.2** | **-18.2** | **15.8** |  | **-11.3** | **-4.6** | **-10.9** |
| **Xinjiang_Marsden** | **-15.9** | **-5.7** | **-7.1** | -0.4 | **5.4** | **3.3** | **15.7** | **7.2** | **4.1** | 1.4 | 2.2 | 0.3 |  | **-9.1** | **-12.3** |
| **Qinghai** | **-18.8** | **-13.7** | **-9.6** | **-9.8** | **3.5** | **3.8** | **-4.0** | **-5.4** | **-3.9** | **-16.5** | **9.6** | **7.3** | **-5.6** |  | **24.8** |
| **Tibet** | **-15.4** | **-13.2** | **-9.5** | **-12.9** | **4.6** | **4.4** | -0.1 | **-3.0** | **-3.1** | **-13.2** | **8.2** | **6.5** | **-2.8** | **30.8** |  |

Table S6. D value and Z-score of *D(Fox, Dhole/Jackal; test, Jackal/Coyote/Red_wolf/Zhejiang)*

| **D(O Jackal;P3 P4)** | | | | | | | | | **D(O Dhole;P3 P4)** | |  |  |  |  |  |  |  |
| --- | --- | --- | --- | --- | --- | --- | --- | --- | --- | --- | --- | --- | --- | --- | --- | --- | --- |
|  | **D value** | | | | **Z score** | | | |  | **D value** | | | | **Z score** | | | |
| **P3/P4** | **Dhole** | **Coyote** | **Red_Wolf** | **Zhejiang** | **Dhole** | **Coyote** | **Red_Wolf** | **Zhejiang** | **P3/P4** | **Jackal** | **Coyote** | **Red_Wolf** | **Zhejiang** | **Jackal** | **Coyote** | **Red_Wolf** | **Zhejiang** |
| **Dhole** |  | 0.62 | 0.62 | **0.53** |  | **100** | **100** | **100** | **Jackal** |  | 0.00 | -0.01 | **-0.11** |  | -1.6 | **-4.5** | **-22.4** |
| **Coyote** | -0.62 |  | 0.02 | **-0.13** | **-100** |  | **8** | **-39** | **Coyote** | 0.00 |  | -0.01 | **-0.11** | 1.6 |  | **-3.8** | **-22.5** |
| **Red_Wolf** | -0.62 | -0.02 |  | **-0.15** | **-100** | **-8** |  | **-44** | **Red_Wolf** | 0.01 | 0.01 |  | **-0.11** | **4.5** | **3.8** |  | **-20.8** |
| **Taimyr** | -0.62 | -0.04 | -0.03 | **-0.19** | **-100** | **-21** | **-14** | **-46** | **Taimyr** | 0.01 | 0.01 | 0.01 | **-0.11** | **4.8** | **4.5** | 2.0 | **-16.7** |
| **Guizhou1** | -0.61 | -0.03 | -0.02 | **-0.25** | **-100** | **-17** | **-10** | **-59** | **W12_Guizhou** | 0.02 | 0.01 | 0.01 | **-0.15** | **6.7** | **6.1** | **3.4** | **-22.1** |
| **Guizhou2** | -0.61 | -0.03 | -0.02 | **-0.21** | **-100** | **-16** | **-10** | **-50** | **W13_Guizhou** | 0.02 | 0.01 | 0.01 | **-0.13** | **6.4** | **5.8** | **3.2** | **-21.2** |
| **Jiangxi** | -0.61 | -0.03 | -0.02 | **-0.20** | **-100** | **-16** | **-10** | **-53** | **W2_Jiangxi** | 0.02 | 0.01 | 0.01 | **-0.11** | **6.8** | **5.8** | **3.0** | **-21.1** |
| **Zhejiang** | -0.53 | **0.13** | **0.15** |  | **-100** | **39** | **44** |  | **W7_Zhejiang** | **0.11** | **0.11** | **0.11** |  | **22.4** | **22.5** | **20.8** |  |
| **Jilin** | -0.61 | -0.03 | -0.02 | **-0.18** | **-100** | **-14** | **-8** | **-40** | **W6_Jilin** | 0.02 | 0.02 | 0.01 | **-0.10** | **7.8** | **7.2** | **4.3** | **-13.8** |
| **Heilongjiang** | -0.61 | -0.03 | -0.02 | **-0.16** | **-100** | **-13** | **-7** | **-34** | **W9_Heilongj** | 0.02 | 0.02 | 0.01 | **-0.10** | **7.9** | **7.4** | **4.7** | **-12.9** |
| **Indian** | -0.61 | -0.04 | -0.02 | **-0.18** | **-100** | **-16** | **-11** | **-52** | **Indian** | 0.01 | 0.01 | 0.01 | **-0.11** | **6.0** | **5.3** | 2.1 | **-20.8** |
| **Iran** | -0.61 | -0.04 | -0.02 | **-0.18** | **-100** | **-18** | **-12** | **-59** | **Iran** | 0.01 | 0.01 | 0.00 | **-0.11** | **6.5** | **5.8** | 2.2 | **-23.1** |
| **Portugal** | -0.61 | -0.04 | -0.02 | **-0.18** | **-100** | **-16** | **-11** | **-48** | **Portugal** | 0.01 | 0.01 | 0.00 | **-0.11** | **5.4** | **4.8** | 1.7 | **-20.8** |
| **Inner_Mongolia** | -0.61 | -0.04 | -0.02 | **-0.19** | **-100** | **-19** | **-12** | **-55** | **Inner_Mongolia** | 0.01 | 0.01 | 0.01 | **-0.11** | **6.4** | **5.7** | 2.5 | **-22.3** |
| **Liaoning** | -0.61 | -0.04 | -0.03 | **-0.19** | **-100** | **-19** | **-12** | **-55** | **Liaoning** | 0.01 | 0.01 | 0.00 | **-0.11** | **5.7** | **4.9** | 1.9 | **-22.4** |
| **Xinjiang** | -0.61 | -0.04 | -0.02 | **-0.19** | **-100** | **-20** | **-14** | **-63** | **Xinjiang** | 0.01 | 0.01 | 0.00 | **-0.11** | **5.8** | **5.1** | 1.4 | **-25.5** |
| **Altai** | -0.61 | -0.04 | -0.03 | **-0.18** | **-100** | **-19** | **-12** | **-54** | **Altai** | 0.01 | 0.01 | 0.00 | **-0.11** | **5.1** | **4.3** | 1.2 | **-21.2** |
| **Chukotka** | -0.61 | -0.04 | -0.02 | **-0.19** | **-100** | **-18** | **-12** | **-53** | **Chukotka** | 0.01 | 0.01 | 0.01 | **-0.11** | **6.1** | **5.4** | 2.3 | **-22.0** |
| **Bryansk** | -0.61 | -0.04 | -0.03 | **-0.18** | **-100** | **-19** | **-12** | **-51** | **Bryansk** | 0.01 | 0.01 | 0.00 | **-0.11** | **5.5** | **4.5** | 1.6 | **-19.8** |
| **Shanxi** | -0.61 | -0.04 | -0.02 | **-0.19** | **-100** | **-19** | **-12** | **-55** | **Shanxi** | 0.01 | 0.01 | 0.00 | **-0.12** | **5.4** | **4.6** | 1.5 | **-23.4** |
| **China** | -0.61 | -0.04 | -0.03 | **-0.20** | **-100** | **-18** | **-11** | **-50** | **China** | 0.01 | 0.01 | 0.00 | **-0.11** | **5.8** | **5.1** | 2.0 | **-21.0** |
| **Xinjiang_Marsden** | -0.61 | -0.04 | -0.02 | **-0.18** | **-100** | **-18** | **-13** | **-59** | **Xinjiang_Marsden** | 0.02 | 0.01 | 0.01 | **-0.11** | **6.9** | **6.1** | **2.8** | **-22.9** |
| **Qinghai** | -0.61 | -0.04 | -0.02 | **-0.19** | **-100** | **-16** | **-11** | **-52** | **Qinghai** | 0.01 | 0.01 | 0.00 | **-0.11** | **6.0** | **5.2** | 2.1 | **-20.4** |
| **Tibet** | -0.61 | -0.03 | -0.02 | **-0.19** | **-100** | **-15** | **-9** | **-49** | **Tibet** | 0.01 | 0.01 | 0.01 | **-0.11** | **5.7** | **4.8** | 2.0 | **-20.9** |
