## Supplementary Methods for "Genomic approaches reveal an endemic sub-population of gray wolves in Southern China"

**STAR * METHODS**

Detailed methods are provided in the online version of this paper and include the following:

- CONTACT FOR RESOURCE SHARING

- EXPERIMENTAL MODEL AND SUBJECT DETAILS

- METHOD DETAILS

- + Extraction

- + Library preparation

- + Sequencing and data processing

- + Genotype calling

- QUANTIFICATION AND STATISTICAL ANALYSIS

- + Phylogeny, maximum likelihood and neighbor joining

- + Outgroup and admixture *f_3_-*statistics

- + D-statistics

- + TreeMix

- + F4-ratio test

- DATA AND SOFTWARE AVAILABILITY

**- CONTACT FOR RESOURCE SHARING**

**- METHOD DETAILS**

- + **Extraction**

Six historical wolf skin samples were collected from two museums: National Zoological Museum of China in Beijing, and Natural History Museum of Zoology in Kunming. The Zhejiang wolf came from Lin’an, Zhejiang province in 1974 and the Jiangxi wolf came from Jiangxi province in May 1974, both of which are located in the downstream region of the Yangtze River of South China. The two Guizhou wolves came from Guizhou province, also in South China. The final two wolves came from Northeast China (Heilongjiang province on Jan 24th, 1957, and Jilin province on Feb 11th, 1956, respectively). All these samples were treated by AS2O3 for storage.

We extracted DNA from these six different skin samples. Each sample was shaved with a sterilized razor blade to remove the fur. For each skin sample, we cut 25 mg into small pieces of size <1 mm^3^, using sterilized scissors between each sample, placing the pieces into a PCR clean 2.0 mL DNA LoBind tube (Eppendorf, cat. No. 30108078). For each sample, we rinsed the pieces in 70% ethanol (Sigma Aldrich, cat. No. E7023). The mixture was vortexed at maximum speed for one minute and then spun at 13,200 rpm in a table centrifuge for one minute. Finally, we removed the resulting supernatant. We repeated these steps three times and let the tube stand for five minutes at 40°C for complete ethanol evaporation. We used the remaining skin sample in each tube to prepare 50 uL of DNA extract per sample, using the DNA extraction method described in Dabney J, et al 2013.

- + **Library preparation**

Thirty-five libraries were produced using a double stranded library preparation protocol [1, 2]. Libraries were all treated with uracil-DNA-glycosylase (UDG) and endonuclease (Endo VIII) to remove characteristic ancient DNA deamination [3]. All 35 libraries were PCR amplified using AccuPrimePfx DNA polymerase (Life Technologies) [4]. Sample-specific indexes were introduced into both the P5 and P7 adaptors during this library amplification to make it possible to distinguish samples from the new libraries from any other library [1]. Library concentrations were determined using a NanoDrop 2000 spectrophotometer and 2100 Agilent.

- + **Sequencing and data processing**

We sequenced the libraries using 2×150 bp reads on an Illumina HiSeq Xten platform. Reads were demultiplexed according to the expected index pairs (Table S1) allowing one mismatch on each pair of reads. The resulting paired reads were then merged into a single read, requiring an overlap of at least 11 bp (with one mismatch allowed), using a modified form of SeqPrep [5], in which higher quality bases (and scores) are used in the overlap region. After stripping adapters, merged reads were aligned as unpaired molecules using BWA (v0.6.1) using samse [6]. Reads were considered duplicates if they had the same start and end positions, and all duplicates were removed, keeping only the read for each set of duplicates with the highest quality bases (Table S1).

- + **Genotype calling**

For all but the Jiangxi sample, we used random allele calling, choosing not to determine heterozygous sites, as the sequencing depths for most individuals are low (~15x-0.15x, Table 1). We applied a filter where we ignored the first and last two base pairs of each fragment, required a base pair quality higher than 20, a fragment length of no less than 30, and mapping quality of no less than 30. The W2_Jiangxi sample was sequenced to 37x (Table 1), a high enough coverage to call heterozygotes confidently. Thus, we applied the software GATK 1.3 (v1.3-14-g348f2b) with the Unified Genotyper parameter to determine diploid calls.

- QUANTIFICATION AND STATISTICAL ANALYSIS

- + **Principal components analysis**

To investigate the relationship of the newly sampled individuals to wolf and dog populations, we calculated pairwise allele-sharing distances [7] among all pairs of wolf and dog populations. We applied a principal components analysis (PCA) to the resulting pairwise distance matrix using SMARTPCA [7].

- + **Phylogeny**

We constructed the phylogenetic relationship of 32 grey wolves (*Canis lupus*), two coyotes (*Canis latrans*), two jackals (*Canis aureus*), one red wolf (*Canis rufus*), one dhole (*Cuon alpinus*), and one Andean fox (*Lycalopex culpaeus*) using the MEGA package [8]. Maximum Likelihood and Neighbor-Joining methods were used and support values for each node were inferred using 1000 rapid bootstrap replicates.

- + ***f_3_-*statistics**

We computed statistics of the form *f_3_(X,Y; Dhole),* which measures the shared genetic drift between populations X and Y since their separation from an outgroup (*Dhole*) [9].

- + **D-statistics**

We used *D*-statistics [10-12] of the form *D(Fox, Test; X, Y****)*** and *D(Fox, X; Test, Y)* to formally test the relationship these samples have with different wolf populations. We also used *D(Fox, Jackal/Dhole; X, Jackal/Coyote/Red Wolf/Zhejiang)* to determine the ancient genetic component in the Zhejiang wolf.

- + **TreeMix**

We applied TreeMix [13] to investigate the relationship between the newly sequenced samples and wolf and dog populations. TreeMix determines population structure using maximum likelihood trees and allows for both population splits and potential gene flow by using genome-wide allele frequency data and a Gaussian approximation of genetic drift. To further investigate how well the tree model fits the data, we visualized the matrix of residuals for the tree model with no admixture. We test trees for zero, one, and two migration events and show results for zero and two migration events.

- + ***F4-ratio test***

In the D-statistic analysis, we observed that the Zhejiang wolf showed patterns indicating ancestry from a canid population that separated from wolves earlier than the dhole separated from wolves. To estimate the proportion of ancestry in the Zhejiang wolf that was contributed by admixture with a population more archaic than the dhole, we use the F4-ratio test, which provides an unbiased estimate of the admixture proportion [14]. Briefly, we use the dhole and fox as the two source populations for the wolves (X), assuming an unrooted tree (Figure S9). Then, we expect the admixture proportion, $\alpha$, to be given by the following equation.

$\alpha=\frac{f_{4}\left( Dhole, Jackal; Coyote, X \right)}{f_{4}\left( Dhole, Jackal; Coyote, Andean Fox \right)}$ ,

**- DATA AND SOFTWARE AVAILABILITY**

The short-read sequencing data generated as part of this study will be made available on the GSA (http://gsa.big.ac.cn/) under accession no. PRJCA001135.
